# Conserved herbivore-induced volatile signalling despite divergent VOC–life-history associations in two locally adapted *Arabidopsis thaliana* populations

**DOI:** 10.64898/2026.06.17.732448

**Authors:** Rodrigo R. Granjel, Lucía Martín-Cacheda, Gregory Röder, Iago Izquierdo-Ferreiro, Andrea Martín-Díaz, F. Xavier Picó

**Affiliations:** Basque Centre for Climate Change (BC3), 48940, Leioa, Spain; Department of Ecology, Environment and Plant Sciences, Stockholm University, SE, 10691, Stockholm, Sweden; Misión Biológica de Galicia (MBG-CSIC), Apartado de correos 28, Pontevedra, Galicia 36080, Spain; Institute of Biology, University of Neuchâtel, Rue Emile-Argand 11, Neuchâtel 2000, Switzerland; Centro Oceanográfico de Vigo, Instituto Español de Oceanografía (IEO, CSIC), Subida a Radio Faro 50, 36390 Vigo, Spain; Departamento de Ecología y Evolución, Estación Biológica de Doñana (EBD), Consejo Superior de Investigaciones Científicas (CSIC), 41092 Sevilla, Spain

**Keywords:** *Arabidopsis thaliana*, volatile organic compounds (VOCs), plant–plant signalling, herbivore-induced defences, intraspecific variation, life-history traits, local adaptation, plant–insect interactions

## Abstract

- Volatile organic compounds (VOCs) mediate plant–plant signalling and may contribute to phenotypic differentiation among populations. However, the extent to which VOC-mediated signalling varies among locally adapted populations, and how VOC traits relate to major fitness-related traits, remain poorly understood.
- We conducted a greenhouse experiment using two genetically and phenologically divergent Iberian populations of *Arabidopsis thaliana*. Plants were exposed to herbivory by *Spodoptera exigua*, after which we quantified herbivore-induced VOC emissions, VOC-mediated signalling effects on neighbouring conspecifics, and relationships between VOC traits, flowering time, and seed germination.
- Herbivory altered VOC composition, but overall VOC profiles remained broadly similar between populations despite strong divergence in life-history strategies, constitutive resistance to herbivory, and genetic structure. In contrast, correlations between VOC traits and fitness-related traits differed between populations and herbivory treatments. Nevertheless, receiver plants from both populations exhibited reduced herbivore damage after exposure to herbivore-induced emitters, indicating conserved VOC-mediated signalling.
- Our results suggest that herbivore-induced volatile signalling may represent a relatively conserved component of plant defence across locally adapted populations. In contrast, relationships between VOC traits and life-history variation may reflect population-specific integration of defence and fitness-related traits.

## Introduction

Chemical traits play a central role in plant-herbivore interactions. Among these, volatile organic compounds (VOCs) are of particular interest because they mediate both within-plant defence responses and information transfer among neighbouring individuals. Plants subjected to herbivory emit characteristic blends of VOCs that can be perceived by conspecific and heterospecific neighbours, enabling plant–plant signalling (Heil & Karban, 2010; Karban *et al*., 2014b). In response, neighbouring plants may prime or induce defensive pathways (Martinez-Medina *et al*., 2016; Karban, 2021), increasing resistance to herbivores and potentially affecting survival and reproductive success (Karban *et al*., 2016; Aratani *et al*., 2023). Consequently, variation in VOC emission and VOC-mediated signalling may influence ecological performance and contribute to phenotypic differentiation within and among populations.

Despite their potential adaptive importance, linking chemical traits to plant fitness remains challenging because it requires quantifying how variation in VOC traits affects survival and reproduction (Karban & Maron, 2002). Nevertheless, growing evidence indicates that VOC traits can vary substantially within and among populations, a prerequisite for VOC trait evolution. For example, some plant species exhibit different chemotypes, characterised by pronounced within- or among-population differences in VOC production and emission (Karban *et al*., 2016, 2024; Clancy *et al*., 2020; Eckert *et al*., 2023), which likely reflect divergent microevolutionary trajectories. Researchers have also attempted to relate variation in VOC emission and VOC-mediated signalling to selective pressures such as environmental gradients and local environmental heterogeneity. However, results have been inconsistent across systems (Wason *et al*., 2013; Karban *et al*., 2016; Aartsma *et al*., 2019).

One way to better understand VOC trait variation and its evolutionary implications is to examine VOC traits in broader genetic and phenotypic contexts. However, studies investigating genetic variation in VOC emissions and VOC-mediated signalling have yielded contrasting results. Although genetic differentiation in VOC traits is commonly detected, the extent to which this variation translates into differences in VOC-mediated plant–plant communication varies among systems (Karban & Shiojiri, 2009; Karban *et al*., 2014a; Moreira *et al*., 2016; Martín-Cacheda *et al*., 2023). These findings suggest that VOC traits may form part of broader integrated phenotypes alongside morphological and life-history traits. Correlations among traits may arise through shared genetic architecture and selection acting on multiple aspects of the phenotype (Jiang & Zhang, 2023). Under this framework, selection acting on major fitness-related traits may indirectly shape variation in herbivore-induced VOC emission and VOC-mediated signalling. Nevertheless, signalling outcomes remain strongly context-dependent, as cue perception can vary with environmental conditions, developmental stage, and the simultaneous detection of multiple chemical cues (Karban & Maron, 2002; Holopainen & Gershenzon, 2010; Karban, 2021).

The primary objective of this study was to quantify among-population genetic variation in VOC emissions and VOC-mediated signalling effects between neighbouring conspecifics. To achieve this goal, while accounting for within-population genetic structure, we focused on two wild, well-characterised Iberian populations of the annual mustard *Arabidopsis thaliana* locally adapted to contrasting environments: one from a low-elevation, warm site, and the other from a high-elevation, cold site (de la Mata *et al*., 2024). These populations differ strongly in life-cycle phenology, with individuals from the low-elevation population predominantly exhibiting a spring-cycling strategy characterised by high seed dormancy and early flowering, whereas individuals from the high-elevation population mainly display facultative winter-cycling with variable seed dormancy and delayed flowering. Such contrasting phenological strategies underlie seasonal adaptation in *A. thaliana* at regional and global scales (Debieu *et al*., 2013; Vidigal *et al*., 2016; Marcer *et al*., 2018; Martínez-Berdeja *et al*., 2020; Exposito-Alonso, 2020). Because adaptation acts on integrated phenotypes rather than on isolated traits (Adams & Collyer, 2019), strong divergence in life-history strategies may also be associated with differentiation in other complex traits, including herbivore-induced VOC emission and VOC-mediated signalling. Under this framework, genetic variation in VOC traits may covary with major fitness-related traits as a consequence of trait correlations shaping phenotypes in populations adapted to contrasting environmental conditions.

To test these ideas, we conducted a greenhouse experiment in which individuals from two locally adapted *A. thaliana* populations were exposed to leaf herbivory by larvae of the generalist insect *Spodoptera exigua* Hübner (Lepidoptera: Noctuidae). We quantified herbivore-induced VOC emissions and VOC-mediated signalling between neighbouring conspecifics using multiple maternal lines per population, allowing us to evaluate population-level differentiation while accounting for underlying intraspecific genetic variation. In addition, we characterised the chemical composition of emitted VOCs to examine patterns of variation in VOC profiles between populations and herbivory treatments. Finally, we tested whether VOC traits covaried with major fitness-related traits associated with local adaptation, namely germination rate (inversely related to seed dormancy) and flowering time. Specifically, we predicted that VOC traits would covary with these life-history traits as part of broader integrated phenotypes shaped by adaptation to contrasting environments. In contrast, because previous studies have reported inconsistent patterns of genetic variation in VOC-mediated communication, we treated population differences in signalling responses as an open question. Together, this approach allowed us to assess how local adaptation, life-history divergence, and genetic variation are associated with herbivore-induced VOC emission and VOC-mediated plant–plant signalling in *A. thaliana*, and to shed light on how VOC trait evolution may be intertwined with life-history evolution in plants.

## Materials and Methods

### Study system

#### Source populations

We selected two populations from the Iberian collection of *Arabidopsis thaliana* natural populations (Picó *et al*., 2008; Méndez-Vigo *et al*., 2011; Castilla *et al*., 2020; de la Mata *et al*., 2024) that differ markedly in their geographical, environmental, and ecological attributes (Fig. 1). The first population, Bonanza (Bon; 36.88 °N, 6.29 °W, 2 m a.s.l., Cádiz province), occurs in a coastal stone pine (*Pinus pinea* L.) forest on sandy soils, whereas the second population, Canencia (Cai; 40.87 °N, 3.76 °W, 1,520 m a.s.l., Madrid province), is located in a montane Scots pine (*Pinus sylvestris* L.) forest (Fig. 1b,c). Although forest cover is similar at both sites (66–68%; de la Mata *et al*., 2024), they experience contrasting climates, with Bon being warmer and drier than Cai (mean minimum and maximum annual temperatures: 12.3 and 24.1 °C vs. 2.4 and 14.9 °C; total annual precipitation: 563.7 vs. 735.6 mm; Fig. 1d).

**Figure 1.**
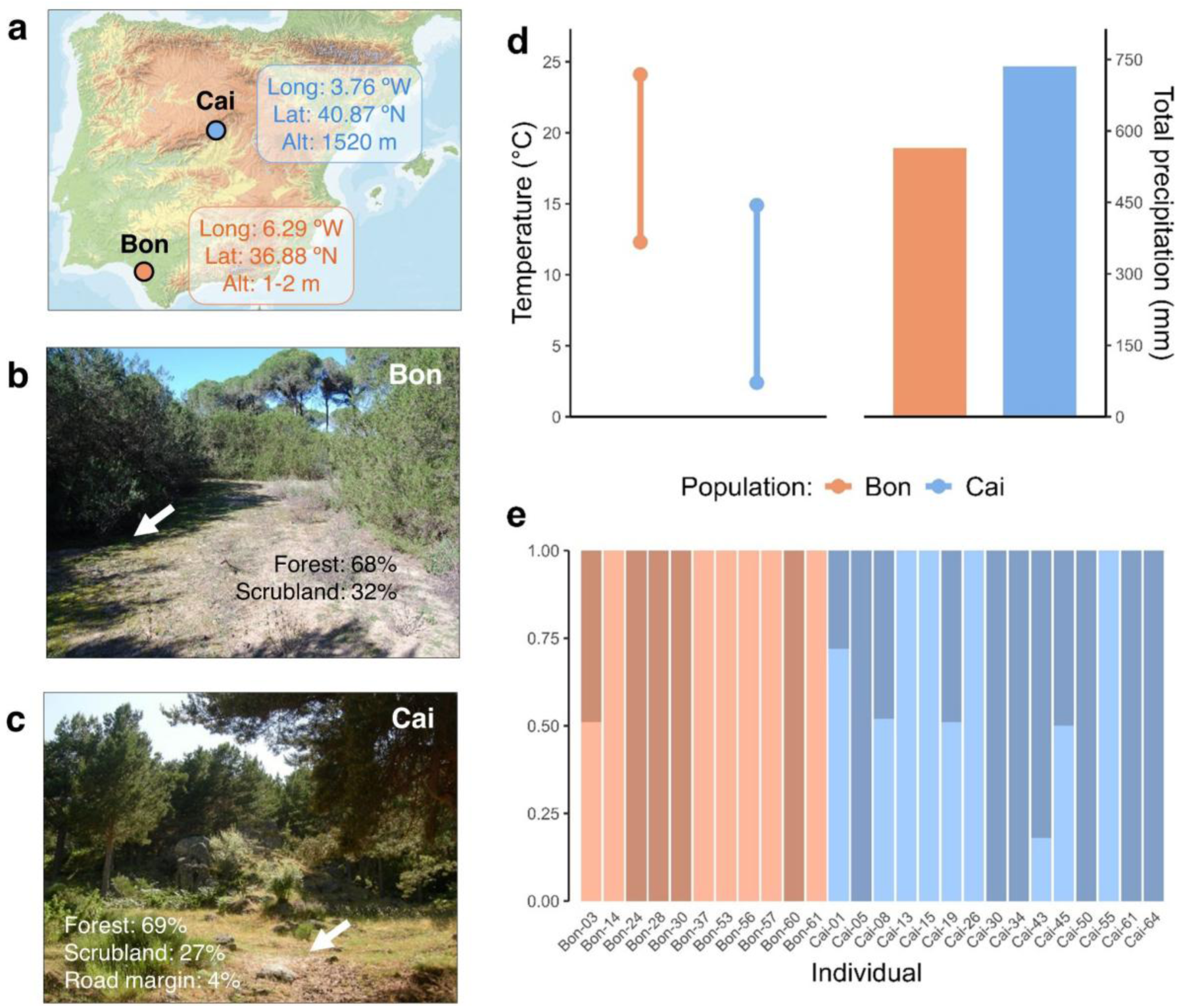
Characterisation of the two *Arabidopsis thaliana* study populations. **(a)** Geographical location of Bonanza (Bon) and Canencia (Cai) in the Iberian Peninsula, with coordinates and elevation. **(b–c)** Photographs of the Bon and Cai sites showing dominant vegetation types and the particular micro-environments where *A. thaliana* occurs within each population (white arrows). **(d)** Climate records for each site, including the average temperature range and the total annual precipitation. (**e**) Genetic cluster membership of individuals from Bon and Cai. Each population comprises two genetic clusters, represented by varying shades of orange (Bon) or blue (Cai).

These climatic contrasts are reflected in population-specific ecological niches. At both sites, *A. thaliana* exhibits a strong preference for forest openings, but plants at Bon occupy shadier and cooler microsites (Fig. 1b), whereas plants at Cai thrive in more exposed and warmer spots (Fig. 1c). Thus, despite sharing a similar forest context, the two populations experience opposite microclimatic optima shaped by their local environments.

Bon and Cai were originally sampled in 2017 as part of a broader study to estimate adaptive life-history variation in Iberian *A. thaliana* (de la Mata *et al*., 2024) and represent the lowest and highest elevations in that dataset. They differ strongly in life-history strategies. At Bon, seeds germinate in mid-winter and plants flower in late winter, shedding seeds in March, consistent with a spring-cycling strategy and prolonged seed-bank dormancy. At Cai, germination spans autumn to late winter, but flowering and seed set occur later in the spring (June), reflecting a mix of winter and spring annual cycles typical of many Iberian natural populations (Montesinos *et al*., 2009; Picó, 2012).

We collected seeds from 50–60 individuals per population and propagated them by single-seed descent in 2018 in a glasshouse at the Centro Nacional de Biotecnología (CNB-CSIC, Madrid, Spain) to minimise maternal and environmental effects. For the experiment, we selected 15 maternal lines per population to maximise spatial and genomic diversity. Mean (±SE) pairwise geographic distances were 74.25 ± 5.65 m at Bon (range: 4.12–146.32 m) and 140.79 ± 8.43 m at Cai (range: 17.00–291.76 m), and mean (±SE) genomic distances (based on allele differences across 2.7 million non-singleton SNPs) were 0.0227 ± 0.0023 at Bon (range: 0.0002–0.0412) and 0.0593 ± 0.0032 (range: 0.0002–0.1029) at Cai. During propagation before the greenhouse experiment (see below), four Bon lines yielded too few viable plants and were discarded, leaving 11 Bon and 15 Cai lines.

Both populations belong to the Iberian relict genetic cluster, an early-diverging lineage associated with long-term climatic stability since the Last Glacial Maximum (Brennan *et al*., 2014; Durvasula *et al*., 2017; Toledo *et al*., 2020; Castilla *et al*., 2020). Despite this shared evolutionary background, Bon and Cai are strongly genetically differentiated and form distinct genetic groups with internal structure (Fig. 1e). Taken together, these differences motivate the use of Bon and Cai as contrasting locally adapted populations for analysing variation in VOC emissions and neighbour-induced responses.

#### Genetic structure

To characterise genetic differentiation between the two study populations, we used a VCF file containing 2,798,036 non-singleton nuclear SNPs from a previous study including 298 maternal lines from six *A. thaliana* populations (de la Mata *et al*., 2024). From this dataset, we extracted the 26 maternal lines used here (11 from Bon and 15 from Cai), yielding 470,456 SNPs. SNP filtering was performed with VCFtools v1.16 (Danecek *et al*., 2011) and PLINK v2.00a3 (Chang *et al*., 2015), retaining loci with <10% missing data and minor allele frequency >5%. We inferred genetic structure using ADMIXTURE v1.3.0 (Alexander *et al*., 2009), which estimates ancestry proportions from SNP genotype data. We used cross-validation and explored K = 1–5 ancestral populations with 20 replicate runs using different random seeds. We selected K = 4, which minimised the cross-validation error (0.4258), and used the resulting ancestry coefficients to visualise genetic structure within and between the two focal populations (Fig. 1e).

#### Herbivore species

We used the generalist chewing herbivore *Spodoptera exigua* Hübner (Lepidoptera: Noctuidae) to induce leaf damage in *A. thaliana*. *Spodoptera exigua* is a highly polyphagous insect that readily feeds on Brassicaceae and is widely used in plant–insect interaction studies (Greenberg *et al*., 2001; Cipollini *et al*., 2004). Furthermore, previous work has shown that leaf damage by *S. exigua* induces strong defensive and volatile responses across a wide range of plant species (Schmelz *et al*., 2003; Danner *et al*., 2018; Martín-Cacheda *et al*., 2023). On this basis, we expected *S. exigua* feeding to induce VOC production and volatile-mediated signalling in *A. thaliana*.

### VOC induction and herbivory experiment

The experiment was conducted at the facilities of the Misión Biológica de Galicia (MBG-CSIC; Salcedo, Pontevedra, Spain). In September 2021, we placed seeds from each maternal line in Petri dishes containing filter paper soaked with deionised water and stratified at 4 °C in complete darkness for four days, followed by five days at 22 °C under continuous light in a FitoClima-10.000-EH growth chamber (ARALAB, Rio de Mouro, Portugal) to promote germination. Germinated seedlings were transplanted with a fine brush into plastic seed trays (104 wells of 40 × 40 × 50 mm^3^; 65 mL per well) filled with a peat-based substrate (Gramoflor GmbH & Co. KG) pre-saturated with water. For each maternal line, we transplanted 48 seedlings to ensure a sufficient number of plants for the experiment.

We maintained trays in a greenhouse under controlled conditions (25 °C during the day and 10 °C at night, with a minimum photoperiod of 10 h). During the first week, we irrigated cells from above every two days and also kept them in shallow water to prevent desiccation; thereafter, we irrigated only from above at the same interval. After four weeks, an average of 27 ± 7 plants per line (57% ± 18% of the original 48 seedlings) reached a rosette diameter of at least 4 cm.

We then separated individual wells from the seed tray and established six pairs of similar-sized plants per maternal line. In each pair, one plant was randomly designated as the emitter and the other as the receiver (Fig. 2). Each emitter-receiver pair was enclosed in a transparent rectangular plastic container (100 × 100 × 220 mm^3^) to ensure isolation and prevent cross-contamination of airborne compounds, resulting in a total of 156 containers. Within each maternal line, three pairs were assigned to the control treatment (no herbivory on emitters) and three to the herbivore-induced treatment (*S. exigua* larvae feeding on emitters). Under the herbivore-induced treatment, one third-instar *S. exigua* larva was placed on each emitter plant, and a nylon mesh was attached to the top of the plant’s well to prevent larval escape. *S. exigua* eggs were provided by the “Control Biotecnológico de Plagas” research group at the University of Valencia, and larvae were reared at the “ECO-EVO” laboratory of the MBG-CSIC.

**Figure 2.**
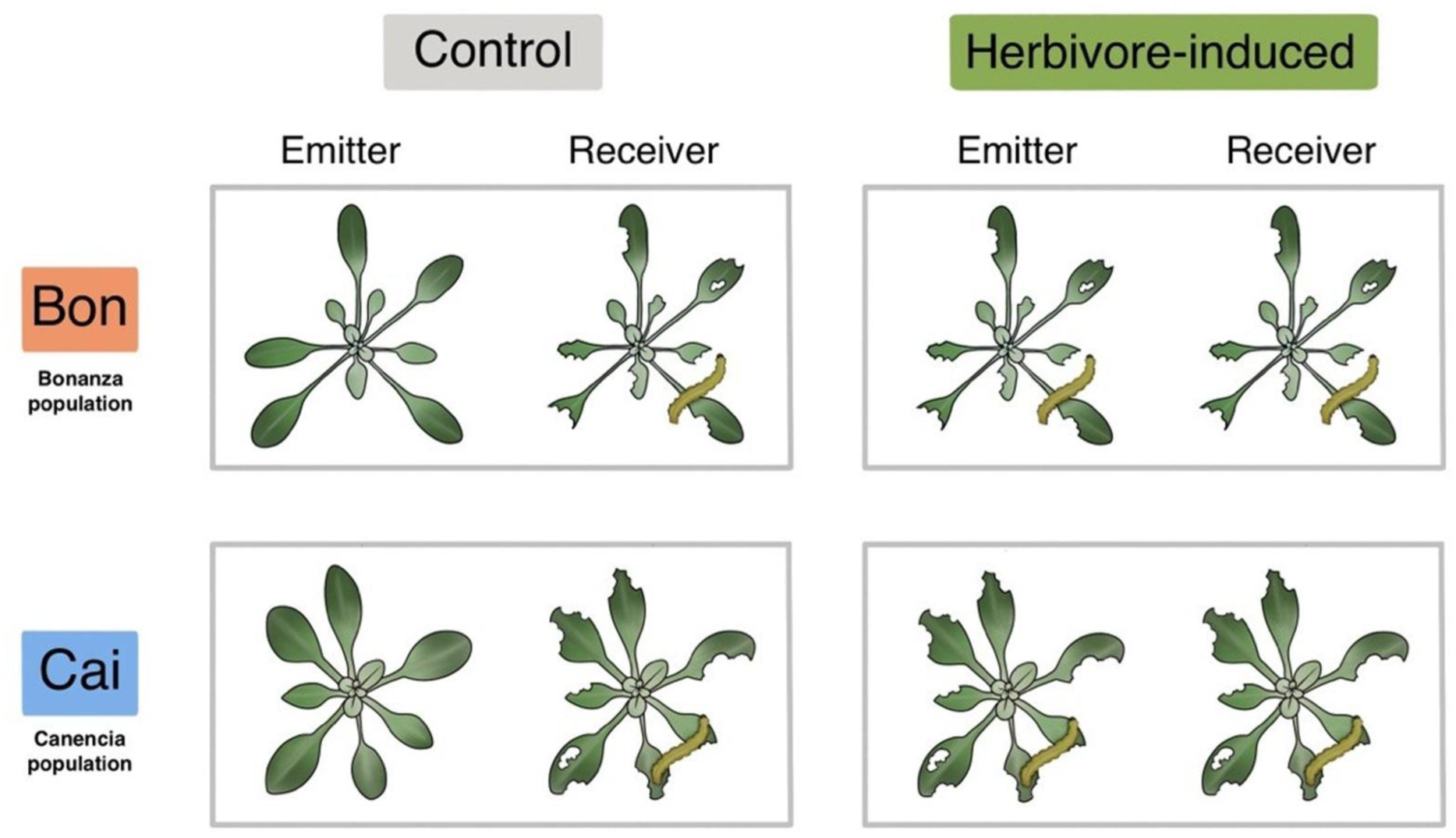
Experimental design. Pairs of *Arabidopsis thaliana* plants from two populations, Bonanza (Bon) and Canencia (Cai), were enclosed in plastic containers (solid boxes) and assigned as emitter or receiver. Emitters were either left undamaged (control) or subjected to herbivory by *Spodoptera exigua* (herbivore-induced). Receivers were subsequently exposed to emitter-derived volatiles and then attacked by caterpillars to quantify feeding damage. This design tests whether volatile organic compound (VOC)-mediated plant–plant signalling alters receiver resistance to herbivores. Population phenotypes are illustrated schematically (Bon: longer, narrower leaves; Cai: broader, more compact leaves). Label colours match those used throughout the manuscript to denote populations (Bon, Cai) and treatments (control, herbivore-induced). Illustration by Rheann Earnest.

Five hours after *S. exigua* feeding, we removed all emitter plants from cages and collected aboveground VOCs from all the emitters following Rasmann *et al*. (2011). To adjust this methodology to the small size of *A. thaliana* plants, we bagged each plant with its container using a 2 L Nalophan bag. We trapped VOCs on a charcoal filter (SKC sorbent tube filled with Anasorb CSC coconut-shell charcoal) for 1.5 h using a Sidekick 224-52MTX pump (SKC Ltd, United Kingdom) (0.25 L min^-1^ airflow of synthetic air N_2_/O_2_). After sampling, charcoal traps were sealed and stored at −80 °C until chemical analysis.

We eluted traps with 150 μL dichloromethane (CAS#75-09-2, Merck, Dietikon, Switzerland) to which we had previously added one internal standard (naphthalene, CAS#91-20-3, 200 ng in 10 μL dichloromethane). We then injected 1.5 μL of the extract for each sample into an Agilent 7890B gas chromatograph (GC) coupled with an Agilent 5977B mass selective detector (Agilent Technologies, Santa Clara, CA, USA) fitted with a 30 m × 0.25 mm × 0.25 μm film thickness HP-5MS ultra inert fused silica column (Agilent Technologies, Santa Clara, CA, USA). We operated the injection into the GC in pulsed splitless mode (250 °C, injection pressure 15 psi) with helium as the carrier gas. The GC oven temperature program was: 3.5 min hold at 40 °C, 5 °C min^-1^ ramp to 230 °C, then a 3 min hold at 250 °C post run (constant helium flow rate 0.9 mL min^-1^). The transfer line was set at 280 °C. In the MS detector (EI mode, 70 eV), a 33–350 (m/z) mass scan range was used with the MS source and the quadrupole set at 230 °C and 150 °C, respectively. We identified volatile compounds using either commercial pure standards or mass spectrum comparisons with the NIST library. For compounds identified by mass spectrum comparison, Kováts indices were calculated relative to the retention times of a series of n-alkanes (C8–C20; Sigma-Aldrich, Merck KGaA, Darmstadt, Germany) analysed under the same chromatographic conditions and compared with values reported in the literature. Although our Kováts indices matched well with previously reported values, compounds not confirmed using commercial standards should be considered putatively identified (Table 1).

**Table 1.** Mean emission rates (ng h^-1^) for individual volatile organic compounds (VOCs) and major VOC classes detected in *Arabidopsis thaliana* emitter plants across populations and treatments. Values in parentheses indicate the 5^th^ and 95^th^ percentiles (P5–P95). Emission rates are expressed as naphthalene-equivalent nanograms per hour based on normalised peak areas. A dagger (†) indicates compounds whose identity was confirmed using commercial standards; unmarked compounds were putatively identified based on mass spectrum comparisons and Kováts indices. Individual compounds are grouped by chemical class (bold text), which broadly reflects different biosynthetic origins: terpenoids derive mainly from the MEP pathway, non-terpenoid aliphatic VOCs from lipid-derived pathways including LOX, and aromatic compounds from the shikimate/phenylpropanoid pathway.

| VOCs | Mean emission (P5–P95) |
| --- | --- |
| **Benzoates** | **1.92 (0.69–4.02)** |
| Acetophenone<sup>†</sup> | 1.92 (0.69–4.02) |
| **Esters** | **2.69 (0.59–7.62)** |
| Butanoic acid, methyl ester | 2.69 (0.59–7.62) |
| **Long-chain aldehydes and alkanes** | **10.11 (0.82–34.44)** |
| Decanal<sup>†</sup> | 15.36 (5.78–45.48) |
| Dodecanal | 18.41 (6.01–45.72) |
| Dodecane<sup>†</sup> | 7.70 (2.34–18.67) |
| Nonanal<sup>†</sup> | 31.13 (11.23–85.95) |
| Tetradecanal | 4.21 (1.41–11.47) |
| Tetradecane<sup>†</sup> | 9.38 (2.20–23.96) |
| Tridecane<sup>†</sup> | 5.85 (1.65–16.51) |
| cis-2-Nonenal | 1.78 (0.47–4.78) |
| **Short-chain oxygenated VOCs** | **9.88 (1.34–27.17)** |
| (E)-2-Methyl-2-butenal | 9.40 (3.38–22.21) |
| 2-Ethyl-1-hexanol<sup>†</sup> | 3.67 (0.96–9.41) |
| 5-Methyl-2-heptanone | 5.57 (2.27–12.14) |
| 6-Methyl-5-hepten-2-one<sup>†</sup> | 24.67 (8.54–72.97) |
| **Terpenoids** | **6.19 (0.71–23.52)** |
| trans-Geranylacetone<sup>†</sup> | 10.70 (2.78–38.65) |
| α-Pinene<sup>†</sup> | 6.30 (1.04–22.94) |
| β-Pinene<sup>†</sup> | 2.17 (0.60–6.02) |

We quantified emissions of individual VOCs using normalised peak areas and expressed them as nanograms per hour (ng h^-1^). For each compound, peak area was divided by the peak area of the internal standard (Abdala-Roberts *et al*., 2022), thereby standardising for variation in sample volume during elution. Reported values for individual VOCs (Table 1; Fig. S1) should thus be considered as naphthalene-equivalent nanograms of compound released per plant per hour. To account for background VOC emissions from soil and containers (Fig. S2), four blank samples were collected under identical conditions as plant measurements. Compound-specific limits of blank (Armbruster & Pry, 2008) were defined as the 95th percentile of blank concentrations for each compound, and signals below this threshold were treated as non-detects in subsequent analyses (61.6% ± 12.6% VOC measurements were retained; Fig. S3). Total VOC emission for each emitter plant was then obtained by summing emissions across individual compounds. For class-level summaries, individual VOCs were grouped according to chemical structure and predominant biosynthetic origin (Rosenkranz & Schnitzler, 2016; Bouwmeester *et al*., 2019), distinguishing terpenoids, which derive mainly from the plastidial MEP (2-C-methyl-D-erythritol 4-phosphate) pathway, non-terpenoid aliphatic VOCs, which derive mainly from lipid-derived pathways including LOX (lipoxygenase), and aromatic compounds, which derive from the shikimate/phenylpropanoid pathway (Grechkin, 1998; Maeda & Dudareva, 2012; Vranová *et al*., 2013; Liu *et al*., 2023). Due to an elution issue, we removed seven VOC samples; therefore, we used 149 samples for statistical analyses (74 control, 75 *S. exigua* feeding). After VOC collection, emitter plants were returned to their containers and left overnight (approximately 15 h) with their paired receivers.

To evaluate whether exposure to VOCs from damaged emitters enhanced resistance to herbivory, and whether this effect differed between populations, we placed one third-instar *S. exigua* larva on each receiver plant using the same procedure applied to emitter plants. Using a Canon EOS 750D and a 55 mm lens (Canon Inc., Tokyo, Japan), we took top-view photographs of all emitter and receiver plants before and after herbivory events by *S. exigua* to visually estimate the percentage of damage (0–100%). To account for potential differences in plant and herbivore size across populations and treatments, we used the pre-herbivory photographs to estimate plant size on a categorical scale from 1 (smallest) to 4 (largest), and we weighed each *S. exigua* larva to the nearest 0.0001 g using an analytical scale (ABS 120-4N; Kern & Sohn GmbH, Balingen, Germany) before placement on the plants.

### Life-history traits

To assess the relationship between life-history traits and VOC emission under control and herbivore-induced treatments in *A. thaliana* populations, we focused on three traits with high adaptive value: seed weight, seed germination, and flowering time. On a separate assay conducted at the experimental facilities of the Estación Biológica de Doñana (EBD-CSIC, Sevilla, Spain) and the Centro Nacional de Biotecnología (CNB-CSIC, Madrid, Spain), we estimated seed weight per maternal line by weighing three batches of 60 filled seeds each, to the nearest 0.1 mg, using a Sartorius BP61S balance (Sartorius AG, Göttingen, Germany). The same three batches were used to estimate the mean seed germination rate per maternal line. Seeds were placed in Petri dishes containing filter paper soaked with deionised water and stratified at 4 °C in complete darkness for four days, followed by five days at 22 °C under continuous light in a FitoClima-10.000-EH growth chamber (ARALAB, Rio de Mouro, Portugal). Germination was scored under a Stemi-2000-C stereomicroscope (Carl Zeiss Optical, Inc., Chester, VA, USA) when the root tip protruded through the seed coat. Flowering time was estimated under greenhouse conditions as the number of days from seedling transplantation to the opening of the first flower, averaging up to 12 full-sibs per maternal line under a 16-hour photoperiod. Seedlings were raised in a growth chamber and transplanted into 28-well trays filled with a 3:1 soil:vermiculite mixture.

### Statistical analyses

All statistical analyses were conducted in R version 4.4.3 (R Core Team, 2025). A full list of the packages used, generated with grateful v. 0.3.0 (Rodriguez-Sanchez & Jackson, 2025), can be found in the Supporting Information.

#### Constitutive resistance to herbivory

To test whether constitutive resistance to *S. exigua* differed between populations, we analysed herbivory damage in emitter plants subjected to herbivory (i.e. the herbivore-induced treatment). Herbivory damage was square-root transformed to improve model assumptions and analysed using a linear model with emitter population (Bon vs. Cai) as the explanatory variable. We evaluated whether larval mass, emitter plant size, and maternal line nested within population improved model fit by comparing models using AIC; however, none of these terms improved the model and were therefore excluded.

#### Effects of population and herbivory on VOC emissions

We analysed total VOC emission as a function of population (Bon vs. Cai), emitter herbivory treatment (control vs. herbivore-induced), and their interaction. We fitted generalised linear mixed-effects models with a Tweedie error distribution and log link, including maternal line nested within population as a random intercept. We evaluated larval mass and emitter plant size as covariates by comparing models using AIC; however, neither improved model fit, and they were not retained in the final model. We used the same fixed- and random-effects structure to analyse treatment- and population-dependent changes in major VOC classes (Table 1), again using Tweedie generalised linear mixed-effects models with a log link.

We tested for differences in VOC composition using a permutational multivariate analysis of variance (PERMANOVA) based on Bray-Curtis dissimilarities of individual compound abundances (Bray & Curtis, 1957; Anderson, 2001), with population, treatment, and their interaction as predictors and 10,000 permutations constrained within maternal line nested within population. We visualised multivariate patterns using principal coordinate analysis (PCoA) of the same Bray-Curtis dissimilarities, implemented via distance-based redundancy analysis with conditioning on the alternative factor (i.e. we visualised population effects after conditioning on treatment, and treatment effects after conditioning on population) (Legendre & Anderson, 1999). We plotted group centroids for each population and treatment (Moreira *et al*., 2021) and identified compounds contributing most strongly to ordination structure using fitted vectors (p ≤ 0.05 and *R^2^* ≥ 0.6).

#### Correlations between VOCs and life-history traits

We estimated Pearson’s correlation coefficients between VOC emission and life-history traits (seed weight, seed germination, and flowering time) separately for each VOC compound (17 individual VOCs and total VOCs), population (Bon and Cai), and treatment (control and herbivore-induced). VOC emissions were averaged per maternal line within each population × treatment combination, and both VOC and trait values were then standardised within each correlation by subtracting their mean and dividing by their standard deviation.

#### Signalling effects on receiver plants

To test for signalling effects on receiver resistance, we analysed the percentage of leaf area consumed by *S. exigua* using linear mixed-effects models. Leaf damage was square-root transformed to improve residual normality and modelled as a function of emitter treatment (control or herbivore-induced), population (Bon or Cai), and their interaction, with maternal line nested within population included as a random intercept. We evaluated larval mass and receiver plant size as covariates by comparing models using AIC; only larval mass improved model fit and was therefore retained in the final model. The treatment × population interaction was of special interest, as it tested whether signalling effects in response to herbivory were contingent on population.

## Results

We detected 17 VOCs in the headspace of *A. thaliana* plants across the experiment and grouped them into five major classes: long-chain aldehydes and alkanes, short-chain oxygenated VOCs, terpenoids, esters, and benzoates (Table 1). In descending order of mean emission, the most abundant classes were long-chain aldehydes and alkanes, short-chain oxygenated VOCs, terpenoids, esters, and benzoates.

### Constitutive resistance and VOC emissions across populations

Constitutive resistance to herbivory by *S. exigua* differed between populations (Fig. 3a). Under the herbivore-induced treatment, emitter plants from Bon experienced significantly higher leaf damage than those from Cai (*t*_1,76_ = 3.33, *p* < 0.01; Table S1), indicating lower constitutive resistance in the low-elevation population. In contrast, overall VOC composition showed little separation between populations. PERMANOVA detected no significant population effect on VOC composition (Table S2), and PCoA revealed substantial overlap between Bon and Cai, with the first two axes together accounting for 43.4% of the variation (27.3% and 16.1%, respectively; Fig. 3b). This variation was most strongly correlated with the compounds dodecanal (*R^2^*= 0.66, *p* = 0.001) and (E)-2-methyl-2-butenal (*R^2^*= 0.63, *p* = 0.001).

**Figure 3.**
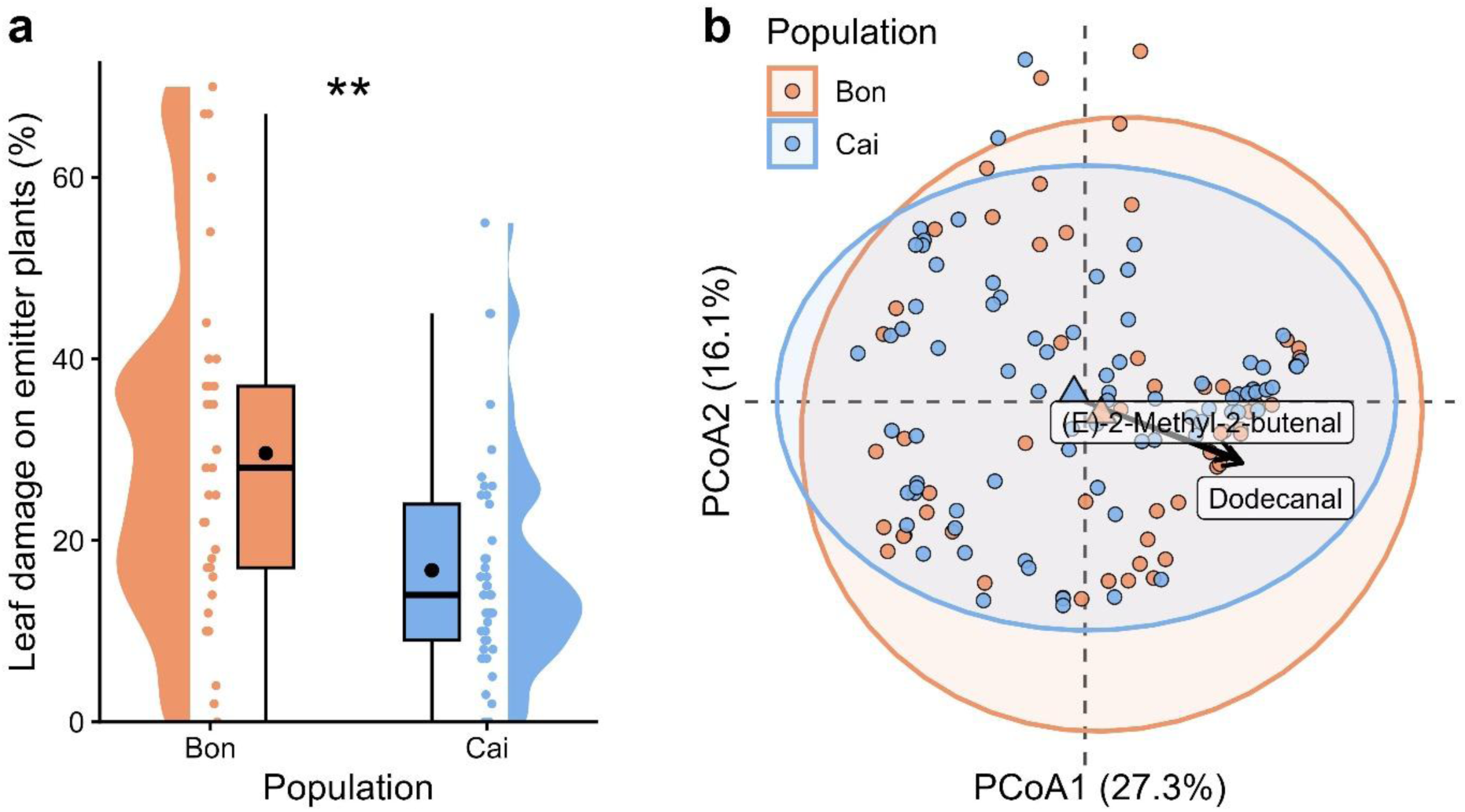
Population-level differences in herbivore damage and volatile organic compound (VOC) emission profiles from emitter plants. **(a)** Percentage of leaf damage caused by *Spodoptera exigua* on emitter plants from two *Arabidopsis thaliana* populations (Bon and Cai). Violin plots show data distribution; boxplots indicate median and interquartile range; dots represent individual observations; black dots indicate means; and asterisks denote statistical significance (\*\**p* < 0.01). **(b)** Principal coordinates analysis (PCoA) of VOCs based on Bray–Curtis dissimilarity. Points correspond to individual plants; arrows and labels represent VOCs significantly correlated with ordination axes; ellipses indicate 95% confidence intervals around population centroids, represented by triangles.

### Effects of herbivory on VOC emissions

Herbivory altered the overall VOC composition across emitter plants. PERMANOVA detected a significant effect of treatment on VOC profiles (*F*_1,127_ = 1.66, *R^2^* = 0.013, *p* = 0.039), as well as a significant treatment × population interaction (*F*_1,127_ = 1.65, *R^2^* = 0.013, *p* = 0.033) (Table S2). Consistent with these results, PCoA based on Bray-Curtis dissimilarities revealed partial separation between control and herbivore-induced emitters, although with substantial overlap between treatments (Fig. 4a). The first two axes together accounted for 43.6% of the variation (27.5% and 16.1%, respectively). Treatment-related variation was most strongly correlated with the compounds tridecane (*R^2^*= 0.61, *p* = 0.001), dodecanal (*R^2^*= 0.65, *p* = 0.001), and (E)-2-methyl-2-butenal (*R^2^*= 0.61, *p* = 0.001).

**Figure 4.**
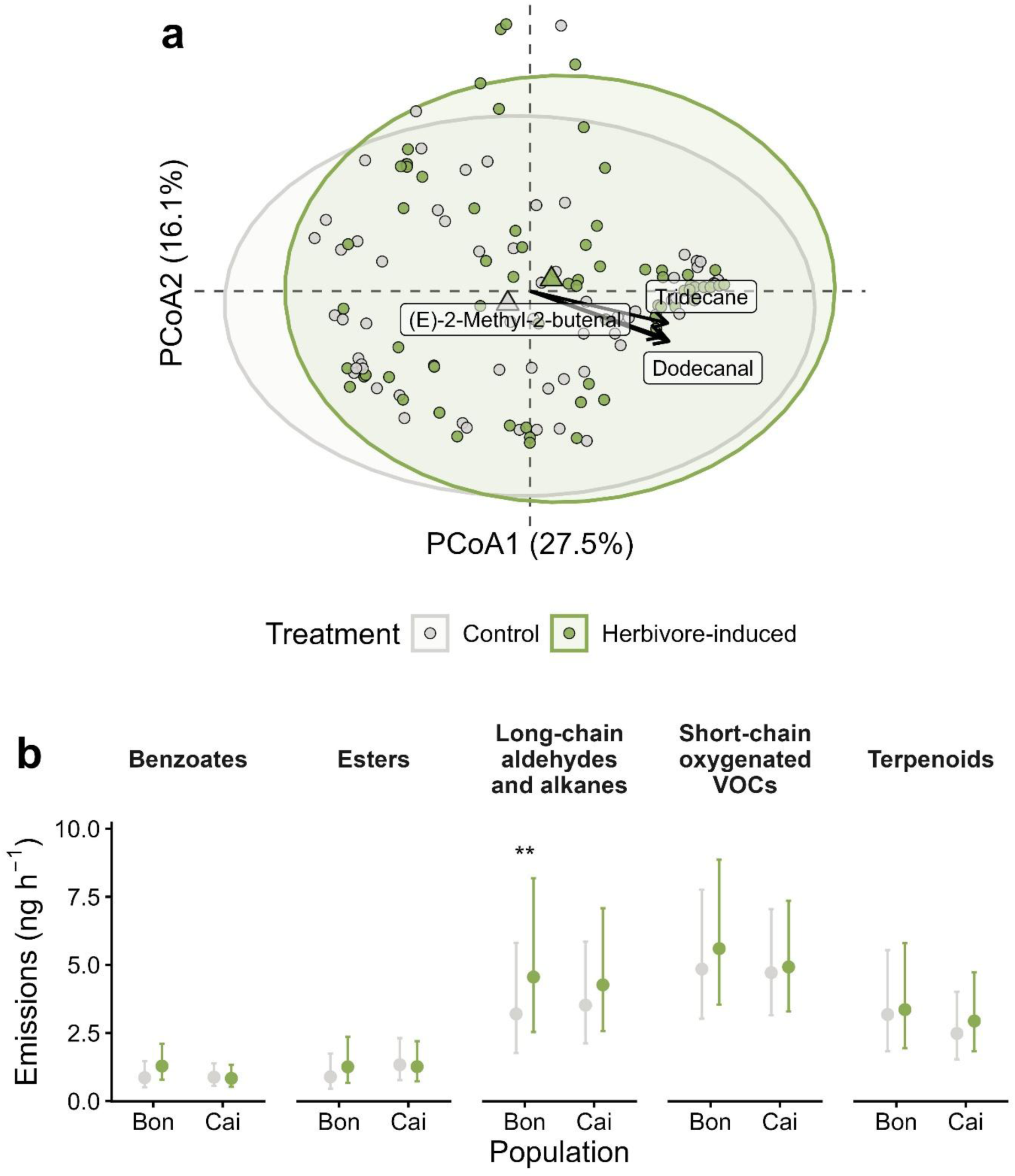
Effects of herbivory on volatile organic compounds (VOCs) emitted by emitter plants. **(a)** Principal coordinates analysis (PCoA) of VOC blends based on Bray-Curtis dissimilarity between *Arabidopsis thaliana* populations (Bon and Cai). Points represent individual plants; arrows and labels indicate VOCs significantly correlated with ordination axes; ellipses show 95% confidence intervals around treatment centroids, represented by triangles. **(b)** Emission rates of the five classes of volatiles analysed. Dots show average emissions, whiskers represent the 95% CI, and asterisks denote statistical significance (\**p* < 0.05; \*\**p* < 0.01; \*\*\**p* < 0.001; no asterisk indicates a non-significant comparison).

At the level of VOC classes, herbivory effects were limited and population dependent (Fig. 4b; Table S3). Emissions of long-chain aldehydes and alkanes increased significantly under herbivory in the Bon population (*z* = −2.85, *p* = 0.004) but not in Cai (*z* = −1.86, *p* = 0.064). No significant treatment effects were detected for benzoates, esters, short-chain oxygenated VOCs, or terpenoids in either population. Total VOC emissions overlapped broadly between control and herbivore-induced treatments in both populations, with no consistent increase under herbivory (Fig. S4; Table S4). Accordingly, herbivory did not significantly affect total emission rates in either population (Bon: *z* = −1.26, *p* = 0.206; Cai: *z* = −0.48, *p* = 0.631).

#### Correlations between VOCs and life-history traits

Correlations between VOC emissions and life-history traits differed between populations and herbivory treatments (Table 2). In Cai, seed germination rate was negatively correlated with several individual constitutive VOCs, whereas no such relationships were observed in Bon. Herbivory maintained these negative correlations in Cai and led to additional compound-specific associations, but the relationship with total VOC emission became non-significant. By contrast, in Bon, several constitutive VOCs were positively correlated with flowering time, whereas no significant correlations were detected in Cai. These relationships disappeared under herbivory, including for total VOC emission. Seed weight showed no significant correlations with any VOC in either population or treatment (all *p* > 0.05).

**Table 2.** Correlation between VOCs and life-history traits. Individual VOCs are grouped by major chemical class (bold headings) and were detected from two Arabidopsis thaliana populations (Bon and Cai) in both control and herbivore-induced treatments. Life-history traits include seed germination and flowering time per population; seed weight showed no significant correlations and is not shown. Pearson’s r-values are given for each VOC–trait correlation, including total VOC emissions. The sample sizes for Bon and Cai were 11 and 15 maternal lines, respectively, across all treatments. Asterisks denote statistical significance (*p < 0.05; **p < 0.01; no asterisk indicates lack of significance). A dagger (†) indicates compounds whose identity was confirmed using commercial standards; unmarked compounds were putatively identified based on mass spectrum comparisons and Kováts indices.

| VOCs | Germination rate |  |  |  | Flowering time |  |  |  |
| --- | --- | --- | --- | --- | --- | --- | --- | --- |
|  | Control |  | Herbivore-induced |  | Control |  | Herbivore-induced |  |
|  | Bon | Cai | Bon | Cai | Bon | Cai | Bon | Cai |
| **Benzoates** |  |  |  |  |  |  |  |  |
| Acetophenone<sup>†</sup> | 0.33 | -0.54 * | 0.20 | -0.56 * | 0.83 ** | 0.05 | 0.46 | -0.01 |
| **Esters** |  |  |  |  |  |  |  |  |
| Butanoic acid, methyl ester | 0.26 | -0.48 | 0.18 | -0.51 | 0.61 * | 0.16 | 0.38 | 0.14 |
| **Long-chain aldehydes and alkanes** |  |  |  |  |  |  |  |  |
| Decanal<sup>†</sup> | 0.21 | -0.69 ** | 0.22 | -0.52 * | 0.61 * | 0.18 | 0.38 | 0.15 |
| Dodecanal | 0.26 | -0.51 | 0.12 | -0.48 | 0.66 * | -0.04 | 0.41 | -0.26 |
| Dodecane<sup>†</sup> | 0.34 | -0.42 | 0.28 | -0.52 * | 0.52 | 0.21 | 0.31 | 0.19 |
| Nonanal<sup>†</sup> | 0.23 | -0.65 ** | 0.30 | -0.64 * | 0.60 | 0.01 | 0.39 | 0.22 |
| Tetradecanal | 0.29 | -0.45 | 0.20 | -0.53 * | 0.68 * | 0.07 | 0.34 | -0.13 |
| Tetradecane<sup>†</sup> | 0.29 | -0.49 | 0.18 | -0.38 | 0.76 * | 0.24 | 0.40 | -0.26 |
| Tridecane<sup>†</sup> | 0.34 | -0.27 | 0.15 | -0.30 | 0.47 | 0.25 | 0.32 | -0.23 |
| cis-2-Nonenal | 0.16 | -0.36 | -0.04 | -0.39 | 0.63 * | 0.15 | 0.31 | 0.20 |
| **Short-chain oxygenated VOCs** |  |  |  |  |  |  |  |  |
| (E)-2-Methyl-2-butenal | 0.21 | -0.42 | 0.02 | -0.47 | 0.78 ** | 0.07 | 0.40 | 0.15 |
| 2-Ethyl-1-hexanol<sup>†</sup> | 0.15 | -0.32 | -0.09 | -0.51 | 0.51 | 0.26 | 0.49 | 0.10 |
| 5-Methyl-2-heptanone | 0.30 | -0.31 | 0.11 | -0.20 | 0.69 * | -0.08 | 0.40 | -0.11 |
| 6-Methyl-5-hepten-2-one<sup>†</sup> | 0.26 | -0.58 * | 0.23 | -0.60 * | 0.66 * | 0.11 | 0.49 | -0.14 |
| **Terpenoids** |  |  |  |  |  |  |  |  |
| trans-Geranylacetone<sup>†</sup> | 0.25 | -0.45 | 0.20 | -0.56 * | 0.68 * | 0.10 | 0.48 | -0.08 |
| α-Pinene<sup>†</sup> | 0.08 | -0.50 | -0.03 | -0.52 * | 0.49 | 0.03 | 0.26 | 0.14 |
| β-Pinene<sup>†</sup> | 0.04 | -0.47 | -0.14 | -0.50 | 0.67 * | 0.10 | 0.43 | -0.11 |
| **Total VOCs** | **0.28** | **-0.60 \*** | **0.19** | **-0.47** | **0.67 \*** | **0.19** | **0.53** | **-0.05** |

### Signalling effects on receiver plants

Emitter treatment significantly affected leaf damage on receiver plants in both populations (Fig. 5; Table S5). In Bon, receivers exposed to control emitters experienced significantly more damage than those exposed to herbivore-induced emitters (*t*_1,127_ = 3.29, *p* = 0.001). A similar pattern was observed in Cai, where damage to receivers was also lower when paired with herbivore-induced emitters (*t*_1,127_ = 2.82, *p* = 0.006).

**Figure 5.**
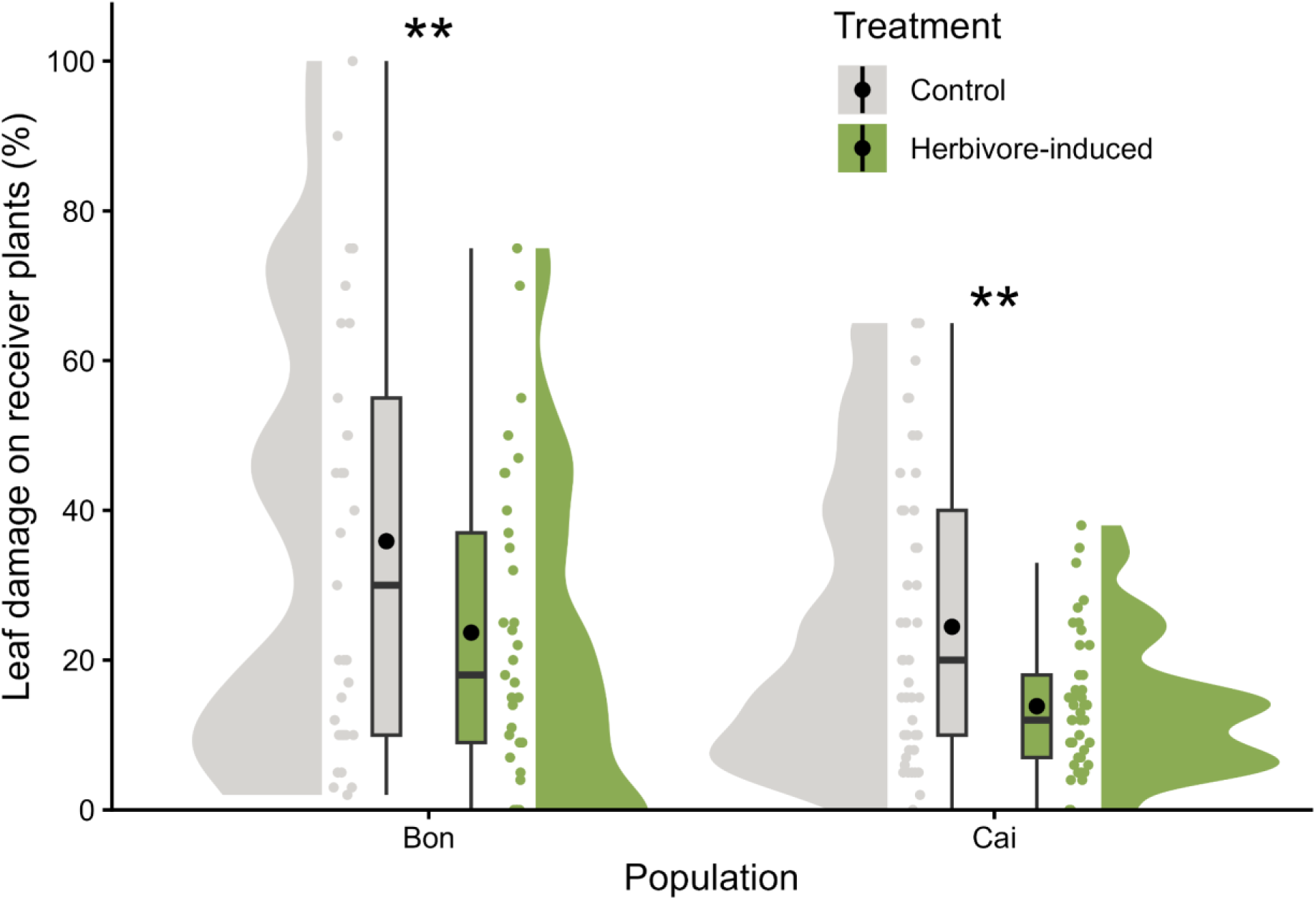
Effects of emitter plant treatment on herbivore damage in receiver plants. Percentage leaf damage caused by *Spodoptera exigua* on receiver plants exposed to control or herbivore-induced emitter plants, shown separately for each *Arabidopsis thaliana* population (Bon and Cai). Violin plots show data distribution; boxplots indicate median and interquartile range; dots represent individual observations; black dots indicate means; and asterisks denote statistical significance (\*\**p* < 0.01).

## Discussion

Herbivore-induced VOCs are important mediators of plant defence and plant–plant communication, yet the extent to which VOC-mediated signalling varies across locally adapted populations remains poorly understood. Because adaptation acts on integrated phenotypes, population divergence in life-history strategies and other fitness-related traits may also influence patterns of VOC emission and VOC-mediated signalling. To investigate these relationships, we used two genetically and phenologically divergent *Arabidopsis thaliana* populations adapted to contrasting environments (Fig. 1) and quantified herbivore-induced VOC emissions, VOC-mediated signalling, and their associations with fitness-related traits. This approach allowed us to examine how local adaptation and life-history divergence relate to variation in inducible volatile-mediated interactions between neighbouring plants.

### VOC-mediated signalling is conserved despite local adaptation

Our results showed that leaf damage by *Spodoptera exigua* on both emitter and receiver *A. thaliana* plants was greater in Bon than in Cai (Figs. 3a, 5), suggesting that plants from Bon were more susceptible or palatable to herbivory than those from Cai. More importantly, receiver plants from both populations experienced lower herbivore damage after exposure to herbivore-induced emitters (Fig. 5), indicating that VOC-mediated signalling was functional and effective in both populations. Herbivore-induced plant–plant signalling has previously been documented in *A. thaliana* (Karban *et al*., 2014b; Estarague *et al*., 2023), although much of the work on volatile-mediated responses in this species has focused on within-plant induced defence responses (Cipollini *et al*., 2004), pathogen-mediated interactions (Riedlmeier *et al*., 2017), and floral scents (Huang *et al*., 2012). In this context, it was particularly notable that herbivore-induced VOC-mediated signalling was conserved despite the marked differences between Bon and Cai in environmental conditions, genetic structure, life-history strategies, and constitutive resistance to herbivory. Together, these findings suggest that herbivore-induced volatile signalling may represent a robust component of plant defence in *A. thaliana*, preserved across broader divergence processes associated with local adaptation.

In terms of VOC composition, constitutive volatile blends were remarkably similar between Bon and Cai (Fig. 3b), and no population-specific VOC chemotypes emerged in our experiment. Herbivory induced weak but detectable shifts in the abundance of some compounds and VOC classes, particularly long-chain aldehydes and alkanes (Fig. 4), although overlap in VOC composition between populations and treatments remained substantial and total emissions did not differ consistently. These results indicate that herbivore-induced volatile blends remained comparatively weakly differentiated despite local adaptation. This degree of differentiation was markedly smaller than the extensive natural variation in VOC profiles previously reported across multiple *A. thaliana* accessions (Duan *et al*., 2005), including phenotypic variation in VOC blends in response to insect herbivory (Snoeren *et al*., 2010; Huang *et al*., 2010). Some of this variation has been linked to allelic differences in genes involved in VOC biosynthesis and defence signalling. For example, Huang *et al*. (2010) showed that differences in herbivory-induced terpene emissions were associated with allelic variation in terpene synthase genes, while Estarague *et al*. (2023) found that genes related to stress-coping ability and jasmonate signalling contributed to variation in snail attraction or repulsion during VOC-mediated plant–plant interactions in *A. thaliana*. By comparison, other defence-related metabolites, such as glucosinolates, exhibit strong geographic structuring and differentiated chemotypes across European *A. thaliana* populations (Katz *et al*., 2021), including contrasting glucosinolate chemotypes in southwestern and central Iberia, where Bon and Cai are located, respectively. Together with our results, these studies suggest that relatively similar VOC blends at the population level may still mask substantial underlying genetic and metabolic variation associated with VOC-mediated interactions in *A. thaliana*.

### Herbivory modifies how VOCs relate to life-history traits across populations

Despite the relatively limited differentiation in VOC composition between populations, relationships between VOC emission and fitness-related traits differed greatly between Bon and Cai. In Bon, early-flowering genotypes emitted higher amounts of VOCs, whereas in Cai, genotypes with stronger seed dormancy (i.e. lower germination rates) also emitted more VOCs (Table 2). These contrasting patterns suggest that different life-history axes structure VOC variation in each population, potentially reflecting population-specific integration between chemical and fitness-related traits. A recent study on the extent of within- and among-population variation in Iberian *A. thaliana*, including Bon and Cai, showed that early flowering was under selection across populations, while traits related to recruitment in Cai, likely including stronger dormancy and reduced germination rates, were also subject to selection (de la Mata *et al*., 2024). Therefore, adaptive differentiation in life-history strategies may indirectly shape VOC-related traits as part of broader integrated phenotypes. More generally, selection acting on recruitment and reproductive schedules in *A. thaliana* (Debieu *et al*., 2013; Marcer *et al*., 2018) may also influence other traits with less direct fitness effects, such as herbivore-induced VOC emissions and VOC-mediated interactions between neighbouring plants.

Herbivory further modified these relationships in contrasting ways between populations. In Bon, the association between flowering time and VOC emission disappeared after herbivore attack, whereas in Cai, the relationship between seed germination and VOC emission persisted under both control and herbivory treatments (Table 2). These contrasting responses suggest that herbivory may alter how VOC-related traits are integrated within broader life-history strategies in each population. Bon and Cai differ strongly in life-cycle dynamics, with spring-cycling genotypes from Bon exhibiting rapid development and early flowering, whereas facultative winter-cycling genotypes from Cai show slower development and delayed reproduction. Under this framework, herbivory may generate different allocation contexts between populations. In Bon, rapid life cycles may favour the maintenance of developmental schedules over sustained coordination between defence-related and life-history traits following herbivore attack, potentially weakening the relationship between flowering time and VOC emission under induced conditions. In contrast, the slower developmental dynamics of Cai may allow VOC-related responses to remain more consistently associated with traits linked to recruitment and dormancy even after herbivore damage. This interpretation is consistent with experimental evidence showing strong genetic integration between vegetative development and flowering initiation in *A. thaliana* through overlapping and pleiotropic quantitative trait loci (Méndez-Vigo *et al*., 2011), as well as broader pleiotropic relationships linking flowering-time genes with multiple developmental and ecological traits (Hanemian *et al*., 2020). Together, these results suggest that herbivory does not simply modify VOC emission, but may also reshape how VOC-related traits are coordinated within locally adapted life-history strategies.

### Perspectives and conclusions

A few aspects of our experimental design should be considered when interpreting these results. Because experiments were conducted under greenhouse conditions using a single herbivore species and a single induction period, patterns of VOC emission and VOC-mediated signalling may differ under more complex natural conditions. In addition, VOC production is known to vary temporally, with some compounds being preferentially emitted during the day and others during the night (De Moraes *et al*., 2001), suggesting that the timing of volatile sampling may influence the composition of detected blends. Our experiment also focused exclusively on rosette-stage plants, whereas flowering *A. thaliana* individuals emit distinct floral volatiles (Huang *et al*., 2012) that could further modify plant–plant interactions and signalling outcomes. Likewise, although we quantified inducible VOCs, we did not measure other defence-related traits, such as constitutive chemical defences, which may also contribute to differences in herbivore resistance between populations.

At the same time, an important strength of our approach is that we incorporated multiple maternal lines per population, allowing us to account for within-population genetic variation rather than relying solely on population-level comparisons. Expanding this framework to additional populations, environmental contexts, developmental stages, and temporal scales may help clarify whether the relationships observed here between VOC emissions and fitness-related traits represent stable components of locally adapted phenotypes or vary according to ecological context and herbivory pressure. Likewise, although our analyses suggest that substantial genetic variation may underlie VOC-mediated interactions, identifying the specific genes and molecular pathways responsible for herbivore-induced VOC production and defence priming remains an important challenge for future research. Future studies could also evaluate the relative importance of direct defence priming through VOC-mediated signalling and indirect defence mechanisms involving the attraction of natural enemies of herbivores (Heil *et al*., 2008). Finally, integrating aboveground and belowground communication pathways, including volatile-mediated signalling and root exudate interactions (Guerrieri & Rasmann, 2024; Subrahmaniam *et al*., 2025), may provide a more complete understanding of how plants coordinate defensive responses to herbivory across heterogeneous environments.

Our study shows that herbivore-induced VOC-mediated signalling remained effective between two locally adapted *Arabidopsis thaliana* populations despite strong divergence in environmental conditions, genetic structure, life-history strategies, and constitutive resistance to herbivory. In contrast, relationships between VOC emissions and fitness-related traits differed markedly between populations and changed under herbivory, suggesting that VOC-related traits may be embedded within population-specific life-history strategies and integrated phenotypes. Together, these findings indicate that herbivore-induced volatile signalling may represent a relatively conserved component of plant defence, while the evolutionary and ecological context in which VOCs are produced and associated with other traits may vary substantially among populations.

## Acknowledgements

We thank Valerie Maas for her assistance during the preliminary experimental phase, Salvador Herrero for providing eggs of *S. exigua*, Beatriz Lago-Núñez for rearing them, and the Ecology and Evolution of Plant-Herbivore Interactions Lab (ECO-EVO, MBG-CSIC) for their support and for providing the facilities necessary for the experiment. We are grateful to Carlos Alonso-Blanco, Belén Méndez-Vigo, and Mercedes Ramiro for their help multiplying field-collected seeds and estimating flowering time under greenhouse conditions at the Centro Nacional de Biotecnología (CNB-CSIC). The Laboratorio de Biología Experimental (LBE) at the Estación Biológica de Doñana and the Doñana Singular Scientific and Technical Infrastructure (ICTS-RBD) provided logistical support. Permission to sample *A. thaliana* at Bon was issued by the Estación Biológica de Doñana (EBD-CSIC) through its Research Coordination Office. No permission was required to sample Cai. We also thank Rheann Earnest for preparing the illustrations used in Fig. 2. This work was supported by María de Maeztu Excellence Unit 2023-2027 Ref. CEX2021-001201-M, funded by MICIU/AEI/10.13039/501100011033; by the Basque Government through the BERC 2026-2029 program; and by the Spanish Association of Terrestrial Ecology (AEET) through a “Tomando la Iniciativa” project grant awarded to Rodrigo R. Granjel.

## Author contributions

RRG secured funding and planned the research with input from FXP, who also provided *Arabidopsis thaliana* seeds. RRG performed experiments with contributions from LM-C. GR and II-F analysed the chemical and morphometric samples, respectively. Data analyses were carried out by RRG and LM-C, and AM-D and FXP performed genetic analyses. RRG, LM-C, AM-D and FXP wrote the manuscript. All authors reviewed and approved the final version of the manuscript.

## Data availability statement

The data and code supporting the findings of this study are openly available in the following GitHub repository: https://github.com/Granjel/CoVOCom.

## Supporting Information

A Supporting Information document accompanies this article, containing the following elements:

● **Table S1**. EMMs and contrasts for leaf damage across populations.
● **Table S2**. PERMANOVA tests between populations, treatments, and both.
● **Table S3**. EMMs and contrasts for all VOC types.
● **Table S4**. EMMs and contrasts for total VOCs across populations and treatments.
● **Table S5**. EMMS and contrasts for leaf damage across populations.
● **Figure S1**. Individual VOCs by population and treatment.
● **Figure S2**. Individual VOCs from blank samples.
● **Figure S3**. VOC samples above the limit of blanks per compound.
● **Figure S4**. Total VOCs by population and treatment.
● **Software citations**. R packages and versions used in the analytical pipeline.

## References

1. Aartsma Y, Leroy B, van der Werf W, Dicke M, Poelman EH, Bianchi FJJA. 2019. Intraspecific variation in herbivore-induced plant volatiles influences the spatial range of plant–parasitoid interactions. Oikos 128: 77–86.

2. Abdala-Roberts L, Vázquez-González C, Rasmann S, Moreira X. 2022. Test of communication between potato plants in response to herbivory by the Colorado potato beetle. Agricultural and Forest Entomology 24: 212–218.

3. Adams DC, Collyer ML. 2019. Phylogenetic Comparative Methods and the Evolution of Multivariate Phenotypes. *Annual Review of Ecology*, Evolution, and Systematics 50: 405–425.

4. Alexander DH, Novembre J, Lange K. 2009. Fast model-based estimation of ancestry in unrelated individuals. Genome Research 19: 1655–1664.

5. Anderson MJ. 2001. A new method for non-parametric multivariate analysis of variance. Austral Ecology 26: 32–46.

6. Aratani Y, Uemura T, Hagihara T, Matsui K, Toyota M. 2023. Green leaf volatile sensory calcium transduction in *Arabidopsis*. Nature Communications 14: 6236.

7. Armbruster DA, Pry T. 2008. Limit of Blank, Limit of Detection and Limit of Quantitation. Clinical Biochemist Reviews 29: S49–S52.

8. Bouwmeester H, Schuurink RC, Bleeker PM, Schiestl F. 2019. The role of volatiles in plant communication. The Plant Journal 100: 892–907.

9. Bray JR, Curtis JT. 1957. An Ordination of the Upland Forest Communities of Southern Wisconsin. Ecological Monographs 27: 325–349.

10. Brennan AC, Méndez-Vigo B, Haddioui A, Martínez-Zapater JM, Picó FX, Alonso-Blanco C. 2014. The genetic structure of *Arabidopsis thaliana* in the south-western Mediterranean range reveals a shared history between North Africa and southern Europe. BMC Plant Biology 14: 17.

11. Castilla AR, Méndez-Vigo B, Marcer A, Martínez-Minaya J, Conesa D, Picó FX, Alonso-Blanco C. 2020. Ecological, genetic and evolutionary drivers of regional genetic differentiation in *Arabidopsis thaliana*. BMC Evolutionary Biology 20: 71.

12. Chang CC, Chow CC, Tellier LC, Vattikuti S, Purcell SM, Lee JJ. 2015. Second-generation PLINK: rising to the challenge of larger and richer datasets. GigaScience 4: s13742-015-0047–8.

13. Cipollini D, Enright S, Traw MB, Bergelson J. 2004. Salicylic acid inhibits jasmonic acid-induced resistance of *Arabidopsis thaliana* to *Spodoptera exigua*. Molecular Ecology 13: 1643–1653.

14. Clancy MV, Haberer G, Jud W, Niederbacher B, Niederbacher S, Senft M, Zytynska SE, Weisser WW, Schnitzler J-P. 2020. Under fire-simultaneous volatilome and transcriptome analysis unravels fine-scale responses of tansy chemotypes to dual herbivore attack. BMC Plant Biology 20: 551.

15. Danecek P, Auton A, Abecasis G, Albers CA, Banks E, DePristo MA, Handsaker RE, Lunter G, Marth GT, Sherry ST, et al. 2011. The variant call format and VCFtools. Bioinformatics 27: 2156–2158.

16. Danner H, Desurmont GA, Cristescu SM, van Dam NM. 2018. Herbivore-induced plant volatiles accurately predict history of coexistence, diet breadth, and feeding mode of herbivores. New Phytologist 220: 726–738.

17. De Moraes CM, Mescher MC, Tumlinson JH. 2001. Caterpillar-induced nocturnal plant volatiles repel conspecific females. Nature 410: 577–580.

18. Debieu M, Tang C, Stich B, Sikosek T, Effgen S, Josephs E, Schmitt J, Nordborg M, Koornneef M, Meaux J de. 2013. Co-Variation between Seed Dormancy, Growth Rate and Flowering Time Changes with Latitude in *Arabidopsis thaliana*. PLOS ONE 8: e61075.

19. Duan H, Huang M-Y, Palacio K, Schuler MA. 2005. Variations in CYP74B2 (Hydroperoxide Lyase) Gene Expression Differentially Affect Hexenal Signaling in the Columbia and Landsberg erecta Ecotypes of Arabidopsis. Plant Physiology 139: 1529–1544.

20. Durvasula A, Fulgione A, Gutaker RM, Alacakaptan SI, Flood PJ, Neto C, Tsuchimatsu T, Burbano HA, Picó FX, Alonso-Blanco C, et al. 2017. African genomes illuminate the early history and transition to selfing in *Arabidopsis thaliana*. Proceedings of the National Academy of Sciences 114: 5213–5218.

21. Eckert S, Eilers EJ, Jakobs R, Anaia RA, Aragam KS, Bloss T, Popp M, Sasidharan R, Schnitzler J-P, Stein F, et al. 2023. Inter-laboratory comparison of plant volatile analyses in the light of intra-specific chemodiversity. Metabolomics 19: 62.

22. Estarague A, Violle C, Vile D, Hany A, Martino T, Moulin P, Vasseur F. 2023. Plant–herbivore interactions: Experimental demonstration of genetic variability in plant–plant signalling. Evolutionary Applications 16: 772–780.

23. Exposito-Alonso M. 2020. Seasonal timing adaptation across the geographic range of Arabidopsis thaliana. Proceedings of the National Academy of Sciences 117: 9665–9667.

24. Grechkin A. 1998. Recent developments in biochemistry of the plant lipoxygenase pathway. Progress in Lipid Research 37: 317–352.

25. Greenberg SM, Sappington TW, Legaspi BC, Liu T-X, Sétamou M. 2001. Feeding and Life History of *Spodoptera exigua* (Lepidoptera: Noctuidae) on Different Host Plants. Annals of the Entomological Society of America 94: 566–575.

26. Guerrieri E, Rasmann S. 2024. Exposing belowground plant communication. Science 384: 272–273.

27. Hanemian M, Vasseur F, Marchadier E, Gilbault E, Bresson J, Gy I, Violle C, Loudet O. 2020. Natural variation at FLM splicing has pleiotropic effects modulating ecological strategies in Arabidopsis thaliana. Nature Communications 11: 4140.

28. Heil M, Karban R. 2010. Explaining evolution of plant communication by airborne signals. Trends in Ecology & Evolution 25: 137–144.

29. Heil M, Lion U, Boland W. 2008. Defense-Inducing Volatiles: In Search of the Active Motif. Journal of Chemical Ecology 34: 601–604.

30. Holopainen JK, Gershenzon J. 2010. Multiple stress factors and the emission of plant VOCs. Trends in Plant Science 15: 176–184.

31. Huang M, Abel C, Sohrabi R, Petri J, Haupt I, Cosimano J, Gershenzon J, Tholl D. 2010. Variation of Herbivore-Induced Volatile Terpenes among Arabidopsis Ecotypes Depends on Allelic Differences and Subcellular Targeting of Two Terpene Synthases, TPS02 and TPS03. Plant Physiology 153: 1293–1310.

32. Huang M, Sanchez-Moreiras AM, Abel C, Sohrabi R, Lee S, Gershenzon J, Tholl D. 2012. The major volatile organic compound emitted from *Arabidopsis thaliana* flowers, the sesquiterpene (E)-β-caryophyllene, is a defense against a bacterial pathogen. New Phytologist 193: 997–1008.

33. Jiang D, Zhang J. 2023. Detecting natural selection in trait-trait coevolution. BMC Ecology and Evolution 23: 50.

34. Karban R. 2021. Plant Communication. Annual Review of Ecology, Evolution, and Systematics 52: 1–24.

35. Karban R, Grof-Tisza P, Blande JD. 2016. CHEMOTYPIC Variation in Volatiles and Herbivory for Sagebrush. Journal of Chemical Ecology 42: 829–840.

36. Karban R, Maron J. 2002. The Fitness Consequences of Interspecific Eavesdropping Between Plants. Ecology 83: 1209–1213.

37. Karban R, Rasheed MU, Huntzinger M, Grof-Tisza P, Blande J. 2024. Alarm calls of sagebrush converge when herbivory is high. Proceedings of the Royal Society B: Biological Sciences 291: 20241513.

38. Karban R, Shiojiri K. 2009. Self-recognition affects plant communication and defense. Ecology Letters 12: 502–506.

39. Karban R, Wetzel WC, Shiojiri K, Ishizaki S, Ramirez SR, Blande JD. 2014a. Deciphering the language of plant communication: volatile chemotypes of sagebrush. New Phytologist 204: 380–385.

40. Karban R, Yang LH, Edwards KF. 2014b. Volatile communication between plants that affects herbivory: a meta-analysis. Ecology Letters 17: 44–52.

41. Katz E, Li J-J, Jaegle B, Ashkenazy H, Abrahams SR, Bagaza C, Holden S, Pires CJ, Angelovici R, Kliebenstein DJ. 2021. Genetic variation, environment and demography intersect to shape *Arabidopsis* defense metabolite variation across Europe (MC Schuman and A Korte, Eds). eLife 10: e67784.

42. Legendre P, Anderson MJ. 1999. Distance-Based Redundancy Analysis: Testing Multispecies Responses in Multifactorial Ecological Experiments. Ecological Monographs 69: 1–24.

43. Liu Y, Singh SK, Pattanaik S, Wang H, Yuan L. 2023. Light regulation of the biosynthesis of phenolics, terpenoids, and alkaloids in plants. Communications Biology 6: 1055.

44. Maeda H, Dudareva N. 2012. The Shikimate Pathway and Aromatic Amino Acid Biosynthesis in Plants. Annual Review of Plant Biology 63: 73–105.

45. Marcer A, Vidigal DS, James PMA, Fortin M-J, Méndez-Vigo B, Hilhorst HWM, Bentsink L, Alonso-Blanco C, Picó FX. 2018. Temperature fine-tunes Mediterranean *Arabidopsis thaliana* life-cycle phenology geographically. Plant Biology 20: 148–156.

46. Martín-Cacheda L, Vázquez-González C, Rasmann S, Röder G, Abdala-Roberts L, Moreira X. 2023. Volatile-Mediated Signalling Between Potato Plants in Response to Insect Herbivory is not Contingent on Soil Nutrients. Journal of Chemical Ecology 49: 507–517.

47. Martínez-Berdeja A, Stitzer MC, Taylor MA, Okada M, Ezcurra E, Runcie DE, Schmitt J. 2020. Functional variants of DOG1 control seed chilling responses and variation in seasonal life-history strategies in *Arabidopsis thaliana*. Proceedings of the National Academy of Sciences 117: 2526–2534.

48. Martinez-Medina A, Flors V, Heil M, Mauch-Mani B, Pieterse CMJ, Pozo MJ, Ton J, Dam NM van, Conrath U. 2016. Recognizing Plant Defense Priming. Trends in Plant Science 21: 818–822.

49. de la Mata R, Mollá-Morales A, Méndez-Vigo B, Torres-Pérez R, Oliveros JC, Gómez R, Marcer A, Castilla AR, Nordborg M, Alonso-Blanco C, et al. 2024. Variation and plasticity in life-history traits and fitness of wild *Arabidopsis thaliana* populations are not related to their genotypic and ecological diversity. BMC Ecology and Evolution 24: 56.

50. Méndez-Vigo B, Picó FX, Ramiro M, Martínez-Zapater JM, Alonso-Blanco C. 2011. Altitudinal and Climatic Adaptation Is Mediated by Flowering Traits and FRI, FLC, and PHYC Genes in *Arabidopsis*. Plant Physiology 157: 1942–1955.

51. Montesinos A, Tonsor SJ, Alonso-Blanco C, Picó FX. 2009. Demographic and Genetic Patterns of Variation among Populations of *Arabidopsis thaliana* from Contrasting Native Environments. PLOS ONE 4: e7213.

52. Moreira X, Granjel RR, de la Fuente M, Fernández-Conradi P, Pasch V, Soengas P, Turlings TCJ, Vázquez-González C, Abdala-Roberts L, Rasmann S. 2021. Apparent inhibition of induced plant volatiles by a fungal pathogen prevents airborne communication between potato plants. Plant, Cell & Environment 44: 1192–1201.

53. Moreira X, Petry WK, Hernández-Cumplido J, Morelon S, Benrey B. 2016. Plant defence responses to volatile alert signals are population-specific. Oikos 125: 950–956.

54. Picó FX. 2012. Demographic fate of *Arabidopsis thaliana* cohorts of autumn- and spring-germinated plants along an altitudinal gradient. Journal of Ecology 100: 1009–1018.

55. Picó FX, Méndez-Vigo B, Martínez-Zapater JM, Alonso-Blanco C. 2008. Natural Genetic Variation of *Arabidopsis thaliana* Is Geographically Structured in the Iberian Peninsula. Genetics 180: 1009–1021.

56. R Core Team. 2025. R: a language and environment for statistical computing. Vienna, Austria: R Foundation for Statistical Computing.

57. Rasmann S, Erwin AC, Halitschke R, Agrawal AA. 2011. Direct and indirect root defences of milkweed (*Asclepias syriaca*): trophic cascades, trade-offs and novel methods for studying subterranean herbivory. Journal of Ecology 99: 16–25.

58. Riedlmeier M, Ghirardo A, Wenig M, Knappe C, Koch K, Georgii E, Dey S, Parker JE, Schnitzler J-P, Vlot AC. 2017. Monoterpenes Support Systemic Acquired Resistance within and between Plants. The Plant Cell 29: 1440–1459.

59. Rodriguez-Sanchez F, Jackson CP. 2025. grateful: Facilitate citation of R packages.

60. Rosenkranz M, Schnitzler J-P. 2016. Plant Volatiles. In: Encyclopedia of Life Sciences. John Wiley & Sons, Ltd, 1–9.

61. Schmelz EA, Alborn HT, Banchio E, Tumlinson JH. 2003. Quantitative relationships between induced jasmonic acid levels and volatile emission in *Zea mays* during *Spodoptera exigua* herbivory. Planta 216: 665–673.

62. Snoeren TAL, Kappers IF, Broekgaarden C, Mumm R, Dicke M, Bouwmeester HJ. 2010. Natural variation in herbivore-induced volatiles in *Arabidopsis thaliana*. Journal of Experimental Botany 61: 3041–3056.

63. Subrahmaniam HJ, Picó FX, Bataillon T, Salomonsen CL, Glasius M, Ehlers BK. 2025. Natural variation in root exudate composition in the genetically structured *Arabidopsis thaliana* in the Iberian Peninsula. New Phytologist 245: 1437–1449.

64. Toledo B, Marcer A, Méndez-Vigo B, Alonso-Blanco C, Picó FX. 2020. An ecological history of the relict genetic lineage of *Arabidopsis thaliana*. Environmental and Experimental Botany 170: 103800.

65. Vidigal DS, Marques ACSS, Willems LAJ, Buijs G, Méndez-Vigo B, Hilhorst HWM, Bentsink L, Picó FX, Alonso-Blanco C. 2016. Altitudinal and climatic associations of seed dormancy and flowering traits evidence adaptation of annual life cycle timing in *Arabidopsis thaliana*. *Plant*, Cell & Environment 39: 1737–1748.

66. Vranová E, Coman D, Gruissem W. 2013. Network Analysis of the MVA and MEP Pathways for Isoprenoid Synthesis. Annual Review of Plant Biology 64: 665–700.

67. Wason EL, Agrawal AA, Hunter MD. 2013. A Genetically-Based Latitudinal Cline in the Emission of Herbivore-Induced Plant Volatile Organic Compounds. Journal of Chemical Ecology 39: 1101–1111.

