## Supporting Information for "Conserved herbivore-induced volatile signalling despite divergent VOC–life-history associations in two locally adapted *Arabidopsis thaliana* populations"

### Supporting tables

**Table S1.** Estimated marginal means (EMMs) and contrasts for leaf damage caused by *Spodoptera exigua* larvae on *Arabidopsis thaliana* emitter plants across populations (Bon and Cai). **a)** EMMs for each population, with back-transformed means, standard error (SE), degrees of freedom (df), and 95% confidence intervals (CI). **b)** Contrasts comparing populations, with estimates, SE, df, *t* values, and p values. Derived from a linear model of square-root-transformed herbivory damage with population as a fixed effect.

**a) EMMs**

| Population | Response | SE | df | 95% CI |
|---|---|---|---|---|
| Bon | 26.02 | 3.01 | 76 | 20.4—32.4 |
| Cai | 14.47 | 1.93 | 76 | 10.9—18.6 |

**b) Contrasts**

| Contrast | Estimate | SE | df | t | p |
|---|---|---|---|---|---|
| Bon - Cai | 1.3 | 0.39 | 76 | 3.33 | 0.001 |

**Table S2.** Results of the PERMANOVA analysis testing for differences in volatile organic compound (VOC) profiles between populations (Bon and Cai), treatments (control and herbivore-induced), and their interaction. The table includes degrees of freedom (Df), sum of squares (SS), $R^2$, $F$ value, and p value for each factor and their interaction.

| | Df | SS | $R^2$ | F | p |
|---|---|---|---|---|---|
| Population | 1 | 0.16 | 0.005 | 0.7 | 0.408 |
| Treatment | 1 | 0.37 | 0.013 | 1.66 | 0.039 |
| Population x Treatment | 1 | 0.36 | 0.013 | 1.65 | 0.033 |
| Residual | 127 | 28.10 | 0.969 | | |
| Total | 130 | 28.99 | 1.000 | | |

**Table S3.** Estimated marginal means (EMMs) and contrasts for volatile organic compound (VOC) emissions from *Arabidopsis thaliana* plants in the experiment, separated by VOC type: **a)** Benzoates, **b)** Esters, **c)** Long-chain aldehydes and alkanes, **d)** Short-chain oxygenated VOCs, and **e)** Terpenoids. For EMMs, the table includes treatment (control and herbivore-induced), population (Bon and Cai), response (back-transformed mean), standard error (SE), degrees of freedom (df), and 95% confidence intervals (CI). For contrasts, the table includes the treatments being compared, population, ratio of back-transformed means, SE, df, *z* value, and p value. Derived from Tweedie generalised linear (mixed) models with a log link, including treatment, population, and their interaction as fixed effects.

**a) Benzoates**

*a.1) EMMs*

| Treatment | Population | Response | SE | df | 95% CI |
|---|---|---|---|---|---|
| Control | Bon | 0.86 | 0.23 | Inf | 0.5—1.5 |
| Herbivore-induced | Bon | 1.29 | 0.32 | Inf | 0.8—2.1 |
| Control | Cai | 0.88 | 0.20 | Inf | 0.6—1.4 |
| Herbivore-induced | Cai | 0.83 | 0.20 | Inf | 0.5—1.3 |

*a.2) Contrasts*

| Contrast | Population | Ratio | SE | df | z | p |
|---|---|---|---|---|---|---|
| Control / (Herbivore-induced) | Bon | 0.67 | 0 | Inf | −1.766 | 0.077 |
| Control / (Herbivore-induced) | Cai | 1.05 | 0 | Inf | 0.224 | 0.823 |

**Table S3.** *(Continued.)*

### b) Esters

*b.1) EMMs*

| Treatment | Population | Response | SE | df | 95% CI |
|---|---|---|---|---|---|
| Control | Bon | 0.89 | 0.31 | Inf | 0.5—1.7 |
| Herbivore-induced | Bon | 1.26 | 0.40 | Inf | 0.7—2.4 |
| Control | Cai | 1.33 | 0.37 | Inf | 0.8—2.3 |
| Herbivore-induced | Cai | 1.26 | 0.36 | Inf | 0.7—2.2 |

*b.2) Contrasts*

| Contrast | Population | Ratio | SE | df | z | p |
|---|---|---|---|---|---|---|
| Control / (Herbivore-induced) | Bon | 0.71 | 0 | Inf | −1.280 | 0.200 |
| Control / (Herbivore-induced) | Cai | 1.05 | 0 | Inf | 0.258 | 0.796 |

### c) Long-chain aldehydes and alkanes

*c.1) EMMs*

| Treatment | Population | Response | SE | df | 95% CI |
|---|---|---|---|---|---|
| Control | Bon | 3.20 | 0.97 | Inf | 1.8—5.8 |
| Herbivore-induced | Bon | 4.55 | 1.36 | Inf | 2.5—8.2 |
| Control | Cai | 3.52 | 0.91 | Inf | 2.1—5.9 |
| Herbivore-induced | Cai | 4.27 | 1.10 | Inf | 2.6—7.1 |

*c.2) Contrasts*

| Contrast | Population | Ratio | SE | df | z | p |
|---|---|---|---|---|---|---|
| Control / (Herbivore-induced) | Bon | 0.70 | 0 | Inf | −2.853 | 0.004 |
| Control / (Herbivore-induced) | Cai | 0.83 | 0 | Inf | −1.855 | 0.064 |

**Table S3.** *(Continued.)*

**d) Short-chain oxygenated VOCs**

*d.1) EMMs*

| Treatment | Population | Response | SE | df | 95% CI |
|---|---|---|---|---|---|
| Control | Bon | 4.85 | 1.16 | Inf | 3.0—7.8 |
| Herbivore-induced | Bon | 5.60 | 1.31 | Inf | 3.5—8.9 |
| Control | Cai | 4.72 | 0.97 | Inf | 3.2—7.1 |
| Herbivore-induced | Cai | 4.92 | 1.01 | Inf | 3.3—7.4 |

*d.2) Contrasts*

| Contrast | Population | Ratio | SE | df | z | p |
|---|---|---|---|---|---|---|
| Control / (Herbivore-induced) | Bon | 0.87 | 0 | Inf | −0.936 | 0.349 |
| Control / (Herbivore-induced) | Cai | 0.96 | 0 | Inf | −0.323 | 0.747 |

**e) Terpenoids**

*e.1) EMMs*

| Treatment | Population | Response | SE | df | 95% CI |
|---|---|---|---|---|---|
| Control | Bon | 3.18 | 0.90 | Inf | 1.8—5.5 |
| Herbivore-induced | Bon | 3.36 | 0.94 | Inf | 1.9—5.8 |
| Control | Cai | 2.48 | 0.61 | Inf | 1.5—4.0 |
| Herbivore-induced | Cai | 2.94 | 0.71 | Inf | 1.8—4.7 |

*e.2) Contrasts*

| Contrast | Population | Ratio | SE | df | z | p |
|---|---|---|---|---|---|---|
| Control / (Herbivore-induced) | Bon | 0.95 | 0 | Inf | −0.310 | 0.757 |
| Control / (Herbivore-induced) | Cai | 0.84 | 0 | Inf | −1.059 | 0.290 |

**Table S4.** Estimated marginal means (EMMs) and contrasts for total volatile organic compound (VOC) emissions from *Arabidopsis thaliana* emitter plants across populations (Bon and Cai) and treatments (control and herbivore-induced). **a)** EMMs for each treatment within population, with back-transformed means, standard error (SE), degrees of freedom (df), and 95% confidence intervals (CI). **b)** Contrasts comparing treatments within each population, with estimates, SE, df, *z* values, and p values. Derived from a Tweedie generalised linear mixed-effects model with a log link, including treatment, population, and their interaction as fixed effects, and maternal line nested within population as a random intercept.

**a) EMMs**

| Treatment | Population | Response | SE | df | 95% CI |
|---|---|---|---|---|---|
| Control | Bon | 56.32 | 16.50 | Inf | 31.7—100.0 |
| Herbivore-induced | Bon | 76.40 | 21.38 | Inf | 44.2—132.2 |
| Control | Cai | 59.45 | 14.72 | Inf | 36.6—96.6 |
| Herbivore-induced | Cai | 65.61 | 16.09 | Inf | 40.6—106.1 |

**b) Contrasts**

| Contrast | Population | Ratio | SE | df | z | p |
|---|---|---|---|---|---|---|
| Control / (Herbivore-induced) | Bon | 0.74 | 0 | Inf | −1.265 | 0.206 |
| Control / (Herbivore-induced) | Cai | 0.91 | 0 | Inf | −0.481 | 0.631 |

**Table S5.** Estimated marginal means (EMMs) and contrasts for leaf damage caused by *Spodoptera exigua* on receiver *Arabidopsis thaliana* plants across populations (Bon and Cai) and emitter treatments (control and herbivore-induced). **a)** EMMs for each treatment within population, with back-transformed means, standard error (SE), degrees of freedom (df), and 95% confidence intervals (CI). **b)** Contrasts comparing treatments within each population, with estimates, SE, df, *t* values, and p values. Derived from a linear mixed-effects model of square-root-transformed herbivory damage, including treatment, population, and their interaction as fixed effects, larval mass as a covariate, and maternal line nested within population as a random intercept.

**a) EMMs**

| Treatment | Population | Response | SE | df | 95% CI |
|---|---|---|---|---|---|
| Control | Bon | 29.05 | 5.49 | 30.91 | 18.9—41.3 |
| Herbivore-induced | Bon | 17.87 | 4.30 | 30.79 | 10.2—27.7 |
| Control | Cai | 19.94 | 3.90 | 31.15 | 12.8—28.7 |
| Herbivore-induced | Cai | 12.86 | 3.14 | 31.47 | 7.2—20.1 |

**b) Contrasts**

| Contrast | Population | Ratio | SE | df | t | p |
|---|---|---|---|---|---|---|
| Control - (Herbivore-induced) | Bon | 1.16 | 0 | 127.06 | 3.291 | 0.001 |
| Control - (Herbivore-induced) | Cai | 0.88 | 0 | 127.57 | 2.816 | 0.006 |

### Supporting figures

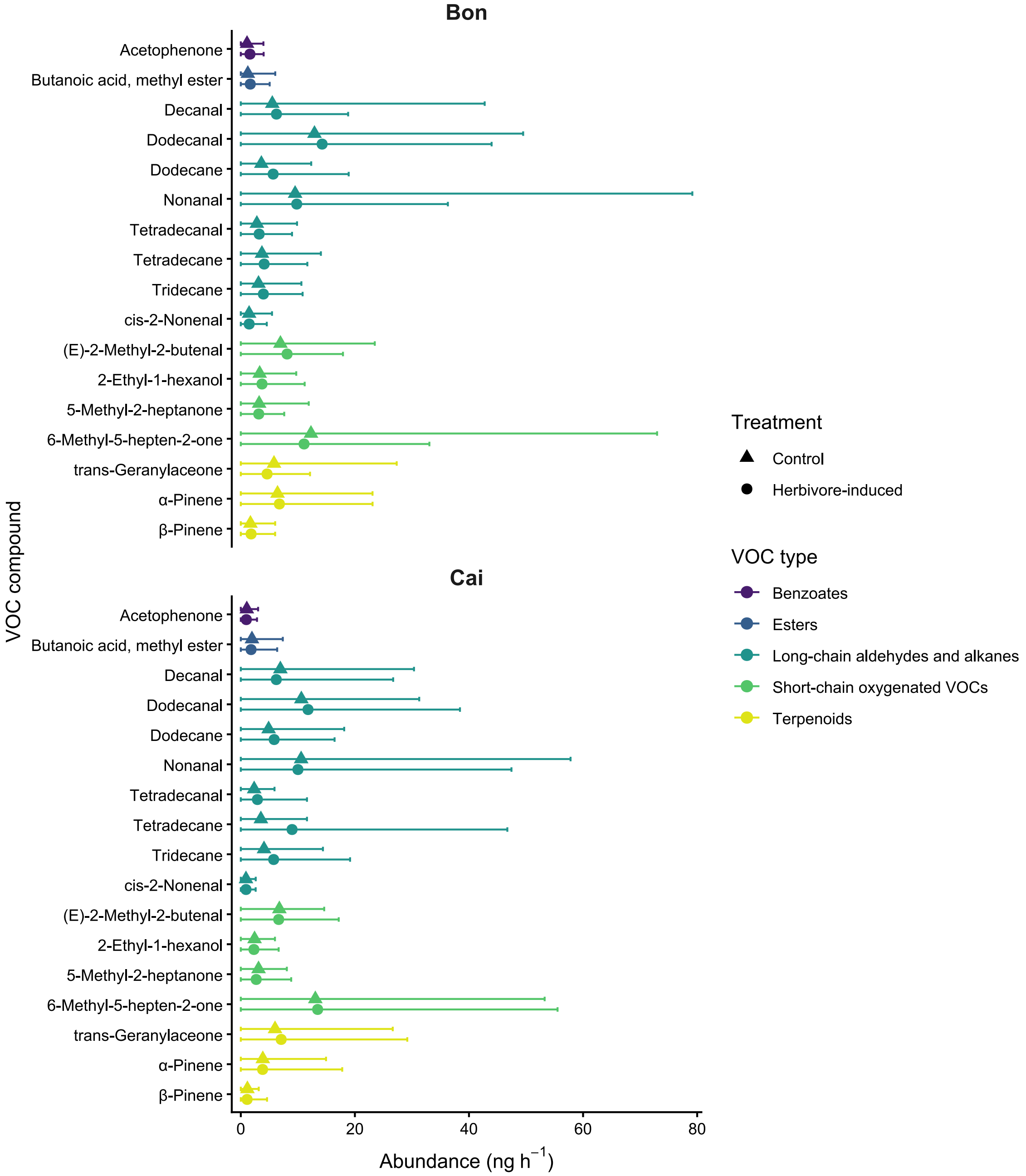


**Figure S1.** Volatile organic compounds (VOCs) from *Arabidopsis thaliana* plants in the experiment, separated by population and treatment. Dots represent mean emissions and error bars represent the 5th and 95th percentiles. Different colors indicate different VOC types.

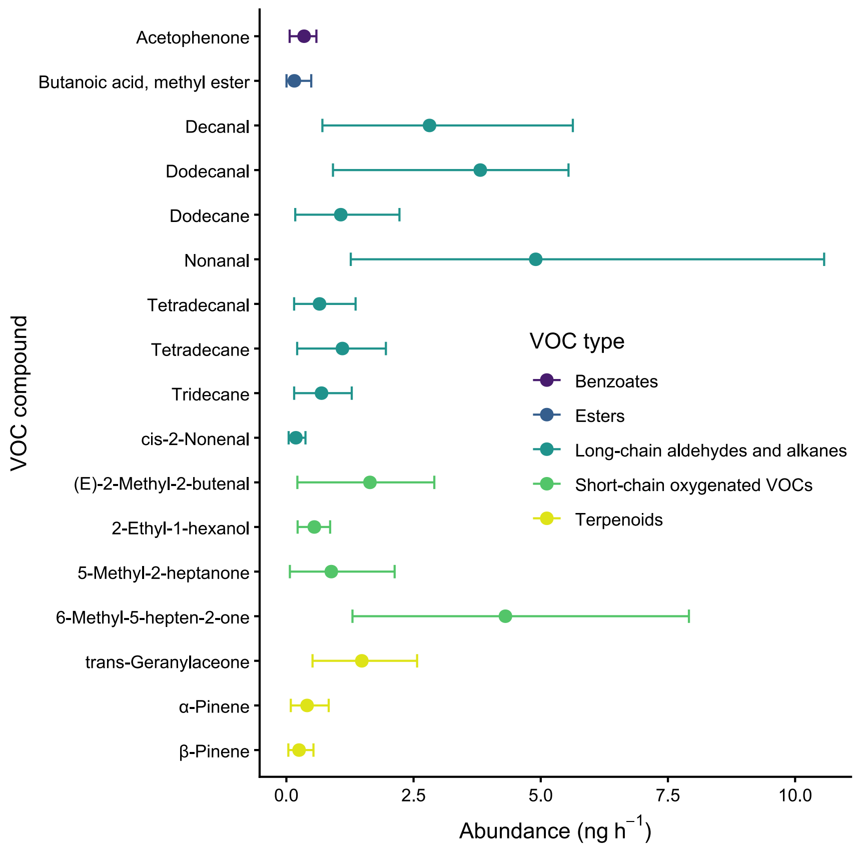


**Figure S2.** Volatile organic compounds (VOCs) from blank samples (no plants, only seedling wells with soil), collected to account for background emissions not produced by plants. Dots represent mean emissions and error bars represent the 5th and 95th percentiles. Different colors indicate different VOC types.

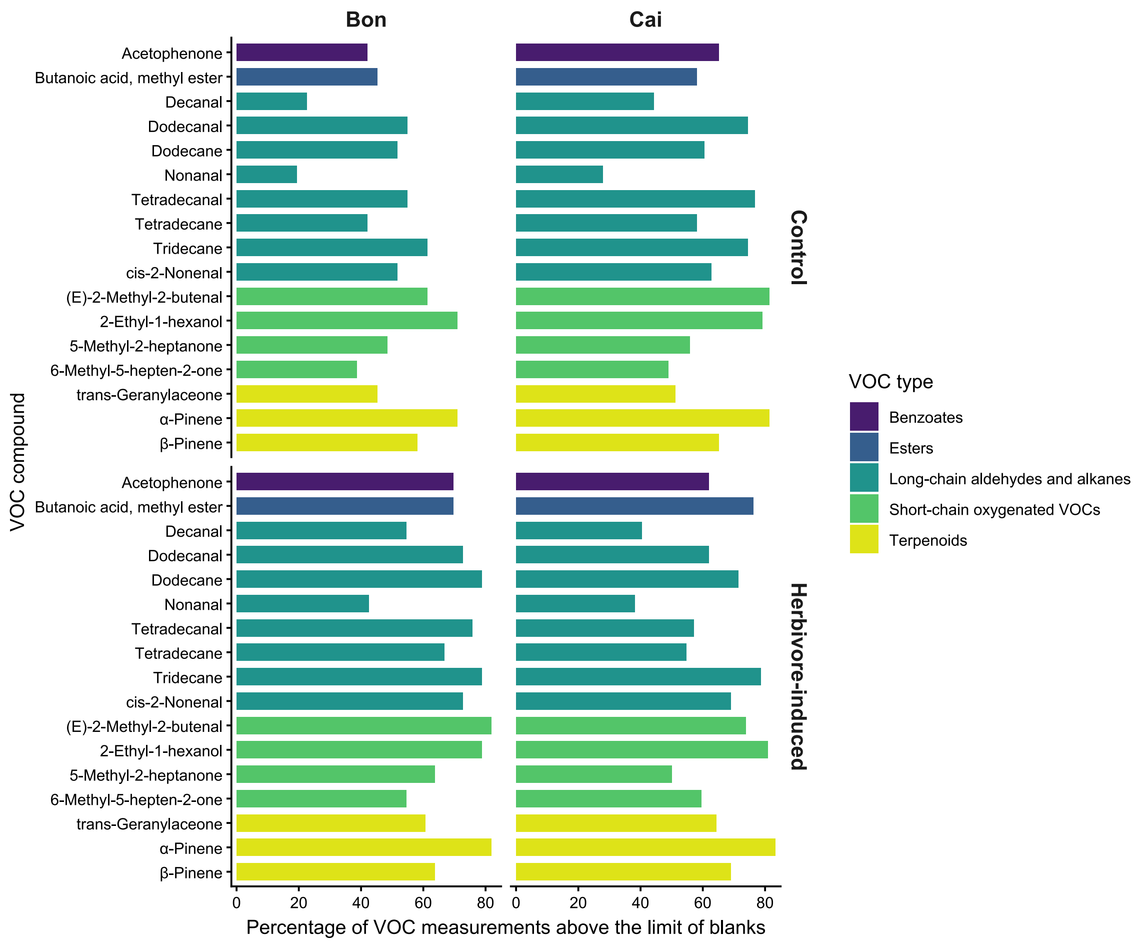


**Figure S3.** Percentage of VOC measurements above the limit of blanks for each compound across maternal lines, for each population and treatment, with the different VOC types indicated by color.

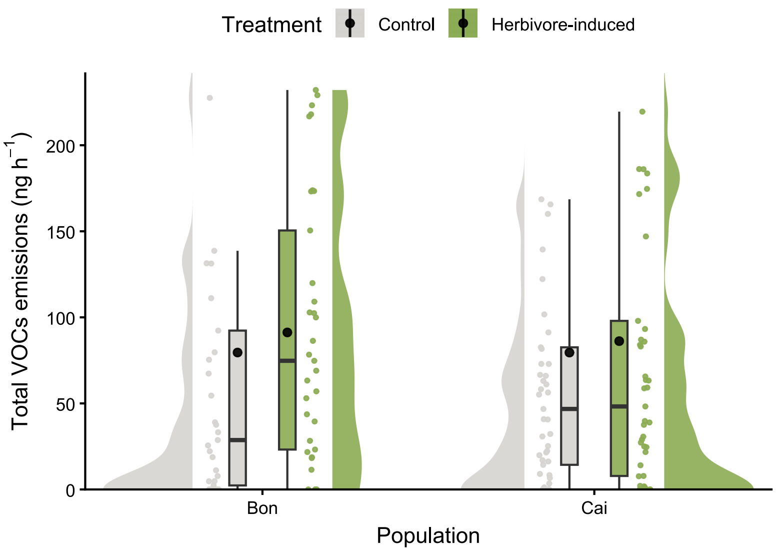


**Figure S4.** Volatile organic compounds (VOCs) from *Arabidopsis thaliana* plants in the experiment, separated by population and treatment. Dots represent mean emissions and error bars represent the 5th and 95th percentiles. Different colors indicate different VOC types.

### Software citations

We used R v. 4.4.3 (R Core Team, 2025) and the following R packages: car v. 3.1.2 (Fox & Weisberg, 2019), carData v. 3.0.5 (Fox *et al.*, 2022), colorspace v. 2.1.1 (Stauffer *et al.*, 2009; Zeileis *et al.*, 2009; 2020), DHARMa v. 0.4.6 (Hartig, 2022), emmeans v. 1.10.1 (Lenth, 2024), ggdist v. 3.3.3 (Kay, 2024; 2025), ggpubr v. 0.6.0 (Kassambara, 2023), ggrepel v. 0.9.5 (Slowikowski, 2024), glmmTMB v. 1.1.9 (Brooks *et al.*, 2017), kableExtra v. 1.4.0 (Zhu, 2024), lattice v. 0.22.6 (Sarkar, 2008), lme4 v. 1.1.35.3 (Bates *et al.*, 2015), lmerTest v. 3.1.3 (Kuznetsova *et al.*, 2017), MASS v. 7.3.64 (Venables & Ripley, 2002), Matrix v. 1.7.2 (Bates *et al.*, 2025), MetBrewer v. 0.2.0 (Mills, 2022), MoMAColors v. 0.0.0.9000 (Mills, 2025), multcomp v. 1.4.25 (Hothorn *et al.*, 2008), multcompView v. 0.1.10 (Graves *et al.*, 2024), mvtnorm v. 1.2.4 (Genz & Bretz, 2009), permute v. 0.9.7 (Simpson, 2022), reshape v. 0.8.9 (Wickham, 2007), survival v. 3.8.3 (Terry M. Therneau & Patricia M. Grambsch, 2000; Therneau, 2024), TH.data v. 1.1.2 (Hothorn, 2023), tidyverse v. 2.0.0 (Wickham *et al.*, 2019), vegan v. 2.6.4 (Oksanen *et al.*, 2022), webshot2 v. 0.1.1 (Chang, 2023).

### Bibliography


**Bates D, Maechler M, Jagan M. 2025.** Matrix: Sparse and Dense Matrix Classes and Methods.

**Bates D, Mächler M, Bolker B, Walker S. 2015.** Fitting Linear Mixed-Effects Models Using lme4. *Journal of Statistical Software* **67**: 1–48.

**Brooks ME, Kristensen K, van Benthem KJ, Magnusson A, Berg CW, Nielsen A, Skaug HJ, Maechler M, Bolker BM. 2017.** glmmTMB Balances Speed and Flexibility Among Packages for Zero-inflated Generalized Linear Mixed Modeling. *The R Journal* **9**: 378–400.

**Chang W. 2023.** webshot2: Take Screenshots of Web Pages.

**Fox J, Weisberg S. 2019.** *An R Companion to Applied Regression*. Thousand Oaks CA: Sage.

**Fox J, Weisberg S, Price B. 2022.** carData: Companion to Applied Regression Data Sets.

**Genz A, Bretz F. 2009.** *Computation of Multivariate Normal and t Probabilities*. Heidelberg: Springer-Verlag.

**Graves S, Piepho H-P, Sundar Dorai-Raj LS with help from. 2024.** multcompView: Visualizations of Paired Comparisons.

**Hartig F. 2022.** DHARMa: Residual Diagnostics for Hierarchical (Multi-Level / Mixed) Regression Models.

**Hothorn T. 2023.** TH.data: TH's Data Archive.

**Hothorn T, Bretz F, Westfall P. 2008.** Simultaneous Inference in General Parametric Models. *Biometrical Journal* **50**: 346–363.

**Kassambara A. 2023.** ggpubr: `ggplot2' Based Publication Ready Plots.

**Kay M. 2024.** ggdist: Visualizations of Distributions and Uncertainty in the Grammar of Graphics. *IEEE Transactions on Visualization and Computer Graphics* **30**: 414–424.

**Kay M. 2025.** ggdist: Visualizations of Distributions and Uncertainty.

**Kuznetsova A, Brockhoff PB, Christensen RHB. 2017.** lmerTest Package: Tests in Linear Mixed Effects Models. *Journal of Statistical Software* **82**: 1–26.

**Lenth RV. 2024.** emmeans: Estimated Marginal Means, aka Least-Squares Means.

**Mills BR. 2022.** MetBrewer: Color Palettes Inspired by Works at the Metropolitan Museum of Art.

**Mills BR. 2025.** MoMAColors: Color Palettes Inspired by Artwork at the Museum of Modern Art in New York City.

**Oksanen J, Simpson GL, Blanchet FG, Kindt R, Legendre P, Minchin PR, O'Hara R, Solymos P, Stevens MHH, Szoecs E, *et al.* 2022.** vegan: Community Ecology Package.

**R Core Team. 2025.** R: A Language and Environment for Statistical Computing.

**Sarkar D. 2008.** *Lattice: Multivariate Data Visualization with R*. New York: Springer.

**Simpson GL. 2022.** permute: Functions for Generating Restricted Permutations of Data.

**Slowikowski K. 2024.** ggrepel: Automatically Position Non-Overlapping Text Labels with `ggplot2'.

**Stauffer R, Mayr GJ, Dabernig M, Zeileis A. 2009.** Somewhere over the Rainbow: How to Make Effective Use of Colors in Meteorological Visualizations. *Bulletin of the American Meteorological Society* **96**: 203–216.

**Terry M. Therneau, Patricia M. Grambsch. 2000.** *Modeling Survival Data: Extending the Cox Model*. New York: Springer.

**Therneau TM. 2024.** A Package for Survival Analysis in R.

**Venables WN, Ripley BD. 2002.** *Modern Applied Statistics with S*. New York: Springer.

**Wickham H. 2007.** Reshaping data with the reshape package. *Journal of Statistical Software* **21**.

**Wickham H, Averick M, Bryan J, Chang W, McGowan LD, François R, Grolemund G, Hayes A, Henry L, Hester J, *et al.* 2019.** Welcome to the tidyverse. *Journal of Open Source Software* **4**: 1686.

**Zeileis A, Fisher JC, Hornik K, Ihaka R, McWhite CD, Murrell P, Stauffer R, Wilke CO. 2020.** colorspace: A Toolbox for Manipulating and Assessing Colors and Palettes. *Journal of Statistical Software* **96**: 1–49.

**Zeileis A, Hornik K, Murrell P. 2009.** Escaping RGBland: Selecting Colors for Statistical Graphics. *Computational Statistics & Data Analysis* **53**: 3259–3270.

**Zhu H. 2024.** kableExtra: Construct Complex Table with `kable' and Pipe Syntax.
